# Behavioral measurements of motor readiness in mice

**DOI:** 10.1101/2023.02.03.527054

**Authors:** Elise N. Mangin, Jian Chen, Jing Lin, Nuo Li

## Abstract

Motor planning facilitates rapid and precise execution of volitional movements. Although motor planning has been classically studied in humans and monkeys, the mouse has become an increasingly popular model system to study neural mechanisms of motor planning. It remains yet untested whether mice and primates share common behavioral features of motor planning. We combined videography and a delayed response task paradigm in an autonomous behavioral system to measure motor planning in non-body-restrained mice. Motor planning resulted in both reaction time savings and increased movement accuracy, replicating classic effects in primates. We found that motor planning was reflected in task-relevant body features. Both the specific actions prepared and the degree of motor readiness could be read out online during motor planning. The online readout further revealed behavioral evidence of simultaneous preparation for multiple actions under uncertain conditions. These results validate the mouse as a model to study motor planning, demonstrate body feature movements as a powerful real-time readout of motor readiness, and offer behavioral evidence that motor planning can be a parallel process that permits rapid selection of multiple prepared actions.

## Introduction

Many goal-directed movements occur too quickly for online correction; the brain plans volitional movements before they are executed. Motor planning has been classically studied in humans and non-human primates using delayed response tasks, in which a temporal delay separates a movement-instructing stimulus from a “Go” cue. These experiments suggest that the delay facilitates a motor planning process after which the brain enters a state of readiness that permits rapid execution of prepared movement (Figure 1A). Motor readiness has been traditionally measured by reaction time (RT, latency to movement onset), which shows that subjects are faster to initiate the movement if given time to prepare. Additionally, the resulting movements are more precise (Figure 1B) (Rosenbaum, 1980; Riehle and Requin, 1989; Churchland et al., 2006b).

**Figure 1:**
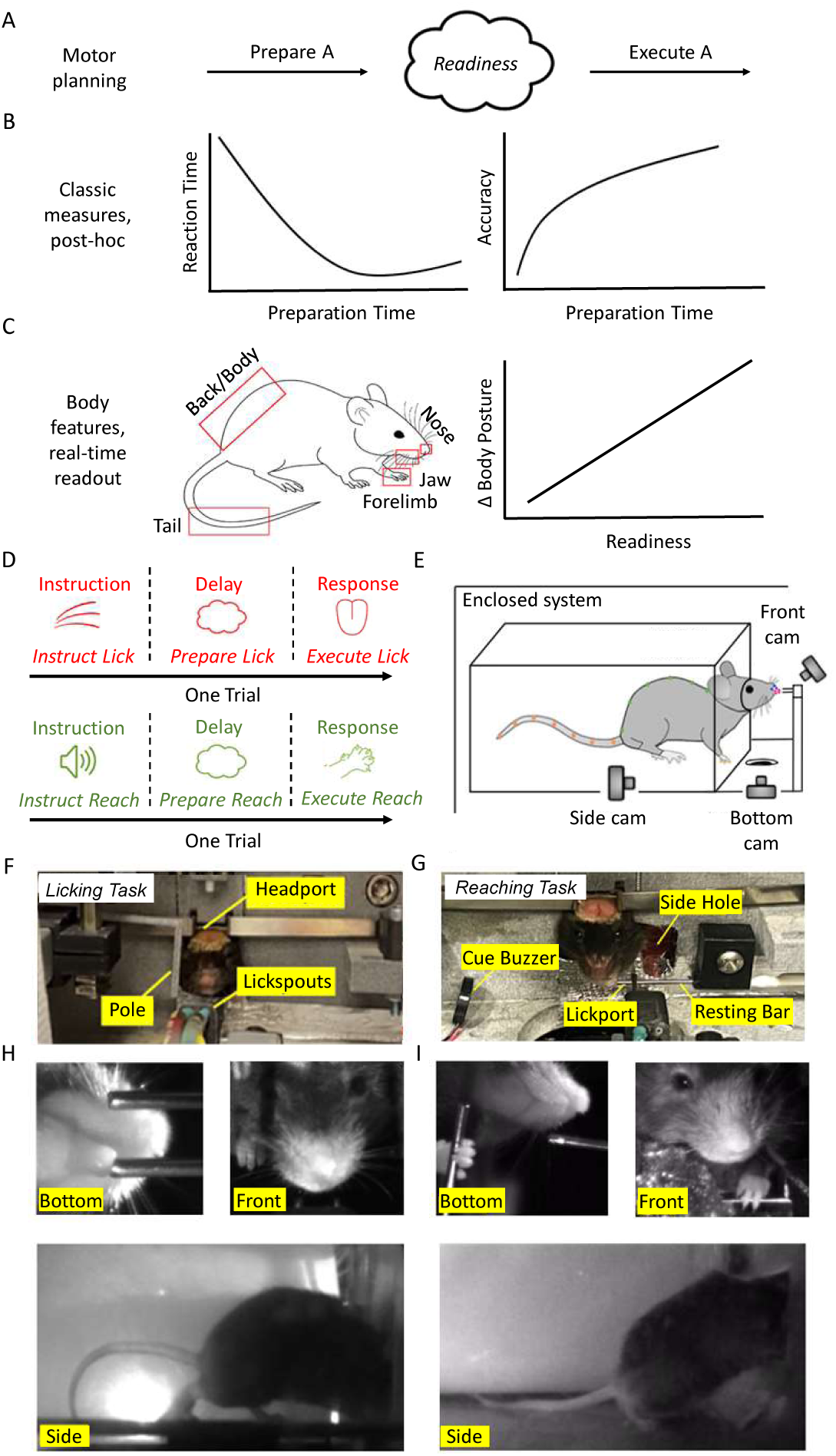
Behavioral measurements of motor readiness, task design and methodology. (A) Cartoon of motor planning. Motor planning leads to a state of motor readiness which allows for rapid and precise movement execution. (B) Classic measurements of motor readiness. *Left*: reaction time decreases with preparation time. *Right*: movement accuracy increases with preparation time. (C) Proposed body feature readout of motor readiness. *Left*: potential body features that reflect motor readiness (jaw, nose, forepaw, back, tail). *Right*: mice might exhibit observable changes in body features during motor planning. (D) Delayed response task paradigm. *Top*: licking task. A tactile cue instructs directional licking after a delay epoch. *Bottom*: reaching task. An auditory cue instructs a reaching movement after a delay epoch. (E) Cartoon of the autonomous behavior system showing location of the front, bottom and side cameras. (F) Front view of the autonomous behavior system configured for the licking task. Image shows a mouse engaged in head-fixation in the headport. (G)Front view of the autonomous behavior system configured for the reaching task. (H) Camera views in the licking task. *Top left*: bottom camera view with the lickport. *Top right*: front camera view. *Bottom*: side camera view. (I) Camera views in the reaching paradigm. *Top left*: bottom camera view with the resting bar and spout. *Top right*: front camera view. Bottom: side camera view.

Many studies have investigated neural substrates of motor planning in non-human primates performing delayed response tasks (Tanji and Evarts, 1976; Riehle and Requin, 1989; Alexander and Crutcher, 1990; Crammond and Kalaska, 2000; Cisek and Kalaska, 2005; Churchland et al., 2010; Michaels et al., 2015). Neurons in motor cortex and connected brain regions show preparatory activity well before movement onset. Preparatory activity has complex temporal patterns and is heterogeneous at the level of individual neurons (Churchland and Shenoy, 2007; Scott, 2008). The prevailing model for these neural dynamics is the dynamical systems framework, which posits that motor planning brings population activity into a specific activity state that is the optimal initial condition for a specific subsequent movement (Afshar et al., 2011; Shenoy et al., 2013). Once entered, activity driving the desired movement can be readily triggered (Churchland et al., 2006b; Kaufman et al., 2014; Gallego et al., 2017). Recently, the mouse has become a popular model system to dissect how neural circuits mediate preparatory activity (Svoboda and Li, 2018). The mouse brain is more amenable to manipulations and comprehensive analyses of neural activity. Preparatory activity has been reported in mouse motor cortex and connected brain regions (Guo et al., 2014a; Li et al., 2015; Guo et al., 2017; Hasegawa et al., 2017; Gao et al., 2018; Chabrol et al., 2019; Chen et al., 2021; Wang et al., 2021). Yet, behavioral measurements of motor planning have thus far been lacking in mice. It remains untested whether motor planning in mice shares the same behavioral features as in humans and monkeys.

Traditional studies of motor planning also have some limitations. Classic delayed response tasks require only one movement to be planned and executed at a time. Correspondingly, the dynamical systems framework posits unique optimal initial conditions for each specific movement. If motor planning is serial (i.e., one movement at a time), it is unclear how the model extrapolates to more naturalistic situations where subjects must select between multiple potential actions, sometimes under time pressure. Recent neurophysiological and behavioral studies suggest that subjects may prepare multiple potential actions simultaneously in uncertain situations and quickly select among them (Cisek and Kalaska, 2010). Neurophysiological recordings in monkeys faced with multiple movement targets show simultaneous representations of potential targets (Cisek and Kalaska, 2005; Cui and Andersen, 2011; Klaes et al., 2011). It remains to be determined whether these activity patterns reflect separate motor plans (Gallivan et al., 2018). Human psychophysics have attempted to measure if subjects simultaneously prepare multiple actions by using probe trials in which the subjects are forced to initiate movements during motor planning (Chapman et al., 2010; Haith et al., 2015; Nashed et al., 2017). However, classic readouts of motor readiness such as movement accuracy and RT are post-hoc measures obtained from movement execution, after an action has been initiated. It remains unresolved whether subjects can simultaneously exhibit motor readiness for multiple actions. Ideally, one would like to read out the state of motor readiness online before movement occurs. An online readout of motor readiness would enable investigations of how motor planning occurs in dynamic situations.

We reasoned that motor readiness might be reflected online in body features during planning (Figure 1C). Volitional movements are often preceded by anticipatory posture changes (Massion, 1992; Dominiak et al., 2019). Moreover, cognitive signals related to decision-making and motor planning could be recorded in downstream motor effectors, such as in spinal cord activity (Prut and Fetz, 1999) and muscle tensions (Selen et al., 2012), in absence of large overt movements. If this is true, we might be able to read out motor planning from body features. Interestingly, recent largescale neurophysiological recordings have also revealed that ongoing posture and movements are reflected in ongoing brain-wide activity (Musall et al., 2019; Stringer et al., 2019; Salkoff et al., 2020; Zagha et al., 2022). Thus, motor planning signals are likely embedded within tightly coupled sensorimotor loops that span the brain and the body. Determining which specific activity contributes to motor planning will be greatly facilitated by real-time measures of motor readiness. Recent advances in video analysis now permit detailed characterization of behavior that could potentially reveal body feature signatures of motor planning (Mathis et al., 2018; Hausmann et al., 2021).

Here, we examined behavioral signatures of motor planning in mice. Non-body-restrained mice voluntarily engaged in two different delayed response tasks in an autonomous behavioral system (Hao et al., 2021): a licking task involving the tongue and a reaching task involving the forelimb. Mice showed RT savings and increased movement accuracy with preparation time, replicating classic effects in humans and monkeys. We found that motor readiness was expressed in preemptive subtle displacements of task-relevant body features, well before movement onset. This behavioral signature followed a time course that tracked the RT savings as a function of preparation time, suggesting a direct relationship to motor readiness. Both the identity of the future action as well as the RT could be predicted from the body feature readout on single trials. We then used this readout to investigate how mice prepare movements under ambiguous situations. Online measures of motor readiness revealed that mice could simultaneously prepare multiple actions in uncertain situations. Together, these results demonstrate that body feature tracking provides a viable real-time readout of motor planning in mice which can be exploited in future studies.

## Results

### Motor planning can be measured in mice using classic measures

To investigate common features of motor planning across different kinds of movements, we developed separate tasks to examine instructed licking and instructed reaching (Figure 1D). In separate sessions, head-fixed mice associated an instructive cue during a sample epoch with either a licking or reaching movement (sample epoch duration: licking task, 1.3s; reaching task, 0.5s) and planned this upcoming movement during a delay epoch (delay epoch duration: both tasks, 1.3s). Mice had to correctly lick or reach the instructed target after an auditory Go cue (pure tone, 3.7kHz, 0.1s duration) to receive a water reward. Mice performed these tasks inside an autonomous behavioral system in which they voluntarily engaged in head-fixation and delayed response tasks by accessing a test chamber through a headport (Hao et al., 2021) (Figure 1E-G). Once head-fixation was initiated, automated computer programs tested mice in the delayed response tasks. In daily sessions, mice were placed into the system for 2 hours in which they learned to perform the task without human supervision (Supplemental Figure 1). In the context of unsupervised testing, we captured mice behaviors using three high-speed cameras (Figure 1H-I; bottom, front and side).

In the licking task, the movement-instructing cue was a tactile stimulus (Guo et al., 2014a). Mice discriminated the position of a vertical pole (anterior vs. posterior) with their whiskers during the sample epoch and reported object location by licking one of two spouts after the delay epoch (lick left or lick right) (Figure 2A). The pole was always presented to the right whisker pad. In the reaching task, the movement-instructing cue was a high-frequency auditory pure tone (10kHz). Mice initiated each trial by using their left forelimbs to hold a horizontal resting bar for 0.5s. The instruction tone played during the sample epoch and mice reached for a drop of water presented on a spout after the delay epoch (Galinanes et al., 2018) (Figure 2B). Mice were not required to maintain the hold on the resting bar after the trial had initiated.

**Figure 2:**
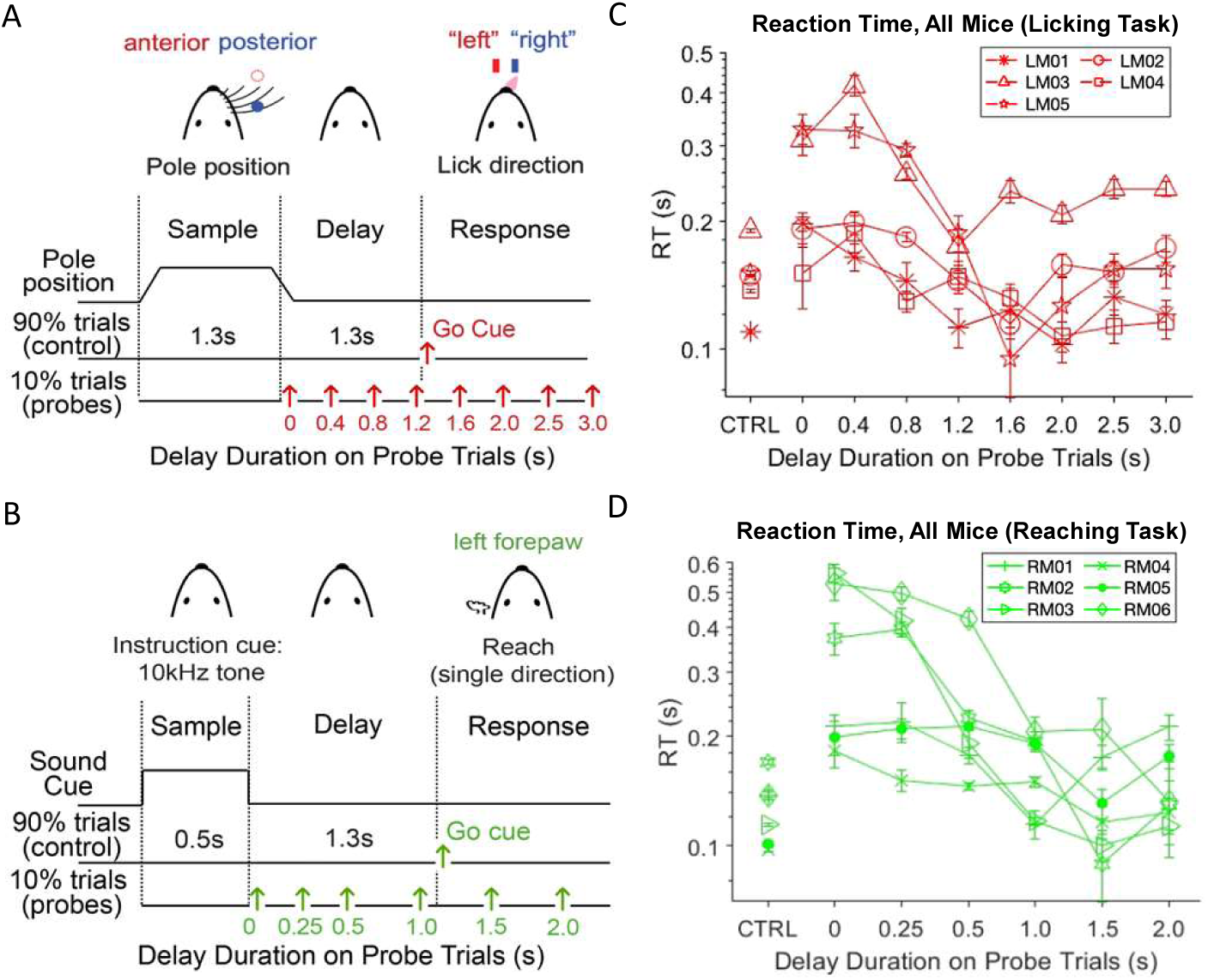
Motor readiness can be measured using classic reaction time measures. (A) Licking task structure. Mice discriminate the location of a pole (anterior or posterior) during a sample epoch (1.3 s) and report the location using directional licking (left or right) after a delay epoch (1.3 s). An auditory Go cue signals the beginning of the response epoch. Red arrows indicate Go cue timing on control trials and the 8 possible probe trial conditions. (B) Reaching task structure. After mice hold a resting bar for 0.5s, a pure tone is played (0.5 s), followed by a delay epoch (1.3 s). An auditory Go cue signals the start of the response epoch. Green arrows indicate Go cue timing on control trials and the 6 possible probe trial conditions. (C) Reaction time (RT) in the licking task for control trials and for each probe trial condition. Individual lines and symbols show the mean and error bars show SEM across trials for individual mouse. (D) Reaction time result in the reaching task. Same as (C).

We first tested whether mice were preparing for specific movements during the delay epoch using classic reaction time (RT) measures. On control trials, the delay epoch spanned 1.3 s. On 10% of trials, we provided the Go cue at an unexpected time (probe trials, Figure 2A-B). Probe trials therefore varied the amount of time animals had to prepare their movement (probe trial delay durations: licking task, 0, 0.4, 0.8, 1.2, 1.6, 2.0, 2.5, 3 s; reaching task, 0, 0.25, 0.5, 1.0, 1.5, 2 s), and they measured the animals’ state of readiness at different time points in a trial. We calculated each mouse’s RTs as the delay between Go cue onset and movement initiation (Methods; Figure 2C-D; control trials, 0.144 ± 0.031 s for the licking task, 0.132 ± 0.023 s for the reaching task, mean ± SD across mice). On probe trials, reaction time was the slowest for the shortest delay durations (Figure 2C-D, 0 s delay duration). RTs decreased with increased delay duration (p<0.05, one-way repeated measure ANOVA test), typically reaching a minimum around the expected time of the Go cue before increasing slightly with additional delay duration (Figure 2C-D). The reaction time savings indicate that mice used the delay epoch to prepare for the upcoming movement, in accordance with classic effects reported in humans and monkeys (Rosenbaum, 1980; Riehle and Requin, 1989; Churchland et al., 2006b; Michaels et al., 2015; Dahan et al., 2019).

Previous studies in primates indicate that movement quality is also an indicator of motor readiness (Riehle and Requin, 1989; Churchland et al., 2006a). In the licking task, mice made more incorrect licks on probe trials with short delay durations compared to long delay durations (Supplemental Figure 2A-B), suggesting reduced movement accuracy with less preparation. To directly examine the impact of motor planning on movement accuracy, we quantified movement trajectory of the first lick or reach following the Go cue. We used DeepLabCut to track the position of the tongue or forelimb (Methods, Figure 3A and F) (Mathis et al., 2018). Because probe trials were limited in number, we examined movement trajectories on control trials. We used RT as a proxy for motor readiness and sorted the trials within each session by RT, yielding a top and bottom percentile (Methods, control trials only). On the slow RT trials, lick and reach trajectories appeared more variable than on the fast RT trials (Figure 3B-C and G-H). To quantify this variability, we calculated the spread of movement endpoints at the target. Movement endpoints were taken as the position of the tongue or forelimb at the end of the protrusion phase and just before the retraction phase. Endpoints were less variable on fast RT trials than slow RT trials (Figure 3D and I, Supplemental Figure 2E). Across all mice, movement endpoints were significantly less variable (more accurate) on fast RT trials than slow RT trials for all trial types and both tasks (Figure 3E and J; Mann-Whitney U Test comparing variance of endpoints in the X and Y dimensions, fast vs. slow RT trials). Similar results were also observed on probe trials where the endpoints were more variable on probe trials with short delay durations than long delay durations (Supplemental Figure 2C-D).

**Figure 3:**
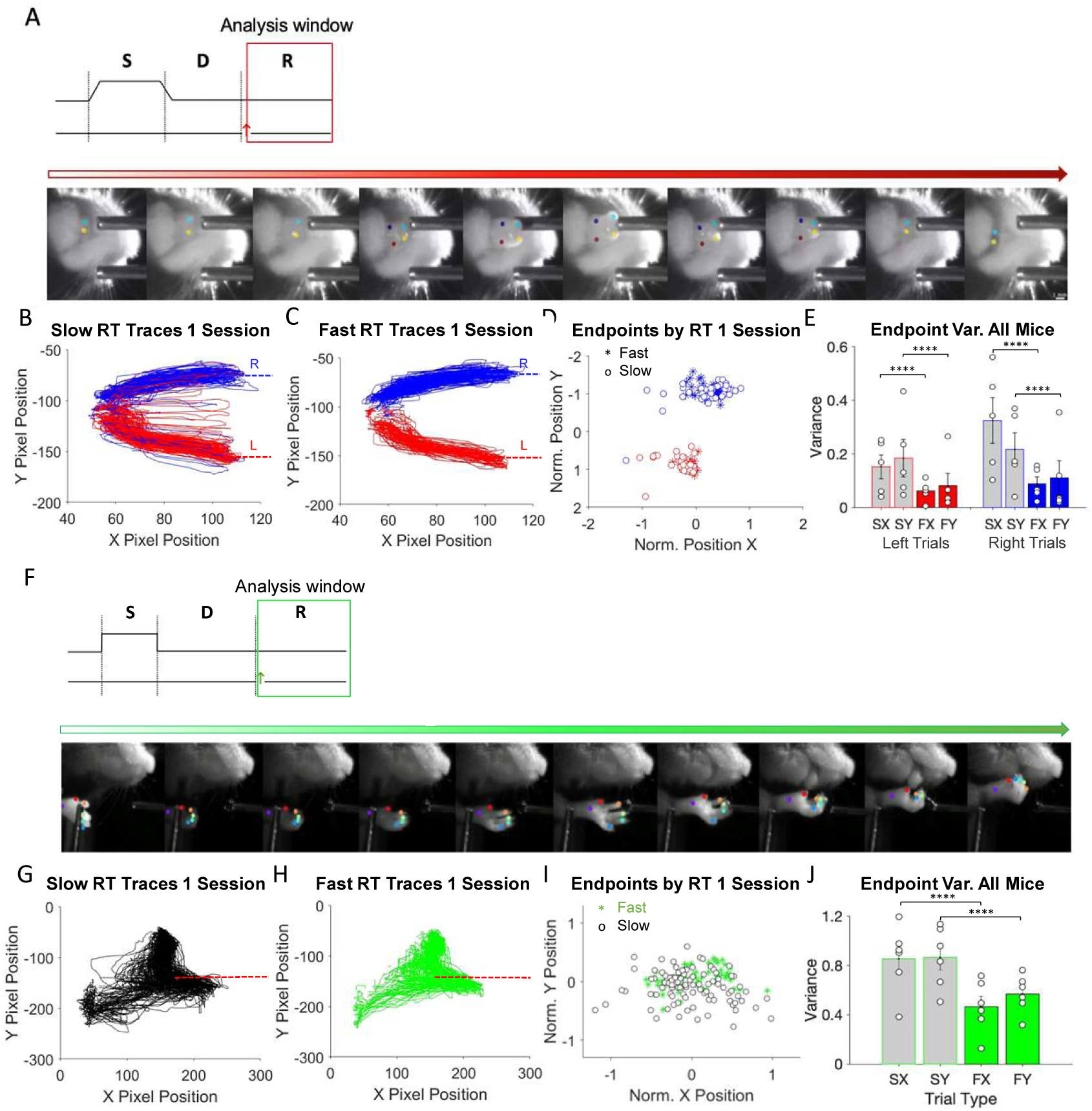
Motor readiness results in more precise movements. (A) *Top*: task timeline and analysis window, the response epoch. *Bottom*: time-lapse frames showing an example tracked lick following the Go cue. (B) Trajectories of the first licks after the Go cue for the slow RT trials in one example session. The slow RT trials are the bottom 1/3 of the trials with the slowest RT from the session. Both correct and incorrect trials are included. Dashed line, lickspout position. (C) Trajectories of the first licks for the fast RT trials. The fast RT trials are the top 1/3 of the trials with the fastest RT from the session. (D) Lick endpoints for the slow and fast RT trials from the same session shown in (B) and (C). Asterisks, fast RT trials; open circles, slow RT trials. (E) Variance of the endpoints on fast RT trials (F) vs slow RT trials (S). Variance are calculated along the X and Y directions for each lick direction (lick left and lick right). Symbols, individual mice; bars, averages; error bars, SD across mice. Lick left trials, X variance, slow vs fast RT trials: p = 0.012, Mann Whitney U two-tailed test; Y variance, slow vs fast RT trials: p = 0.047. Lick right trials, X variance, slow vs fast RT trials: p = 0.036; Y variance, slow vs fast RT trials: p = 0.753. (F)-(J) Same as (A)-(E) for the reaching task. Reach endpoints, X variance, slow vs fast RT trials: p = 0.009, Mann Whitney U two-tailed test; Y variance, slow vs fast RT trials: p = 0.041.

These results show that mice used the delay epoch to prepare for the upcoming movement in the instructed licking and reaching tasks. Motor planning resulted in faster movement initiation and more precise movement.

### Motor planning is reflected online in task-relevant body features

In the real world, sights of motor planning are often visible. For example, baseball pitchers prepare to throw fastballs by positioning their bodies and throwing arms. We tested whether motor planning is reflected in the body features of mice during the delay epoch. We tracked several body features, including the jaw, nose, forelimb, back, and tail. Again using RT during the response epoch as a proxy for motor readiness, we compared the average position of these body features on fast RT trials (more prepared) versus slow RT trials (less prepared; Methods, control trials only). We focused on the position of body features at the end of the delay epoch, averaging across the last five video frames in the delay epoch (Figure 4A and F). A baseline position was also calculated at the start of the delay epoch by averaging across the first five video frames in the delay epoch.

**Figure 4:**
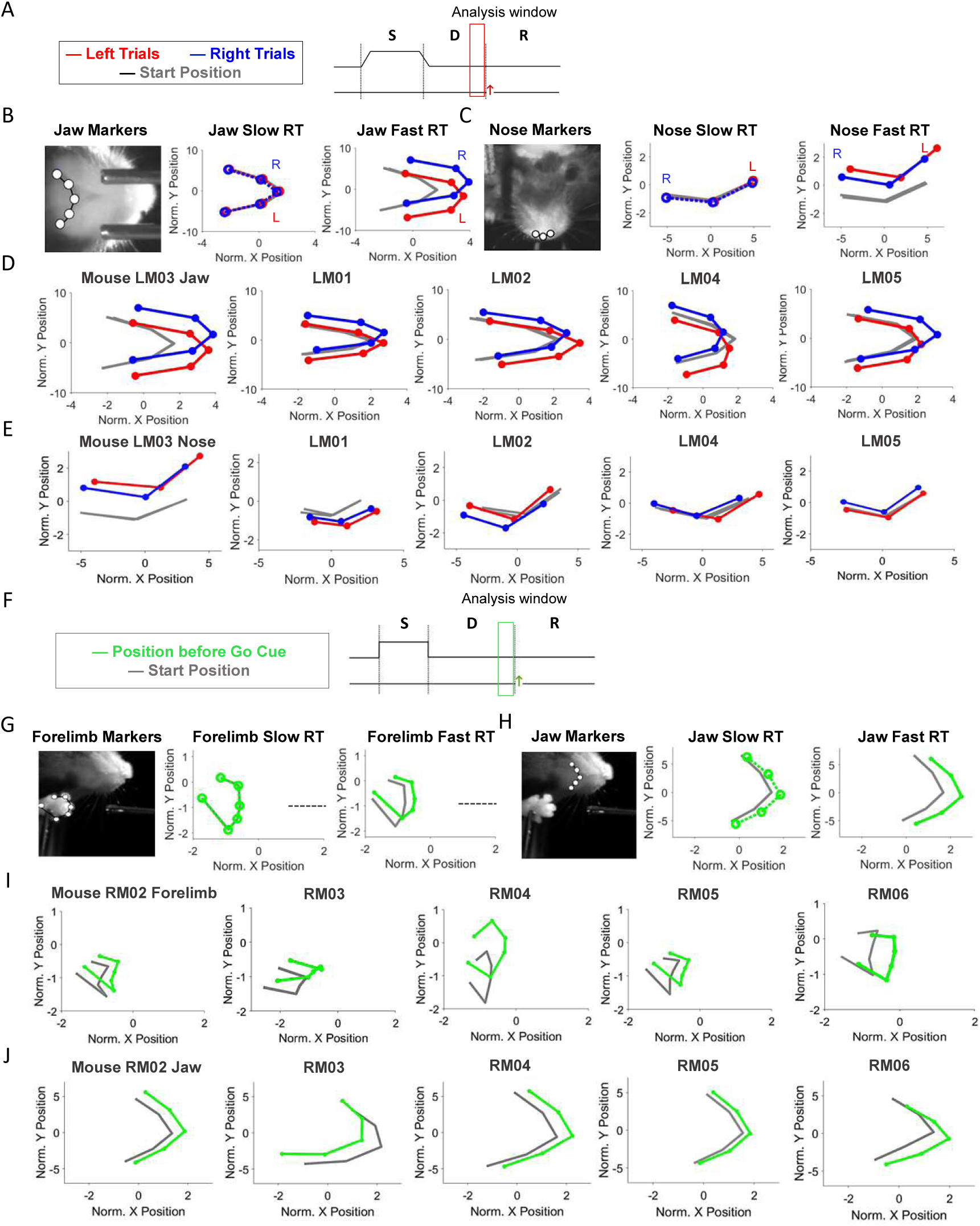
Motor readiness is reflected online in task-relevant body features. (A) Licking task timeline and analysis window, end of the delay epoch. (B) Jaw position from an example mouse. *Left*, location of the five jaw markers. *Right*, the average position of the jaw at the end of the delay epoch on slow and fast RT trials. Red, jaw position on lick left trials; blue, lick right trials. Gray, the average position of the jaw at the start of the delay epoch (baseline). Jaw positions are averaged across all sessions. Only correct trials are plotted. The fast RT trials are the top 1/3 of the trials with the fastest RT from the session. The slow RT trials are the bottom 1/3 of the trials with the slowest RT from the session. (C) Same as (B) for nose position. (D) Jaw positions from all mice. The average position of the jaw at the end of the delay epoch on fast RT trials. Jaw positions are averaged across all sessions for each mouse. (E) Same as (D) for nose position. (F) Reaching task timeline and analysis window, end of the delay epoch. (G)Forelimb position from an example mouse. *Left*, location of the six forelimb markers. *Right*, the averaged position of the forelimb at the end of the delay epoch on slow and fast RT trials. Gray, the average position of the forelimb at the start of the delay epoch (baseline). Spout position is represented by the gray dashed line. Forelimb positions were averaged across all sessions. Only correct trials are plotted. (H) Same as (G) for jaw position. (I) Forelimb position for all mice. The average position of the forelimb at the end of the delay epoch on fast RT trials. Forelimb positions were averaged across all sessions for each mouse. (J) Same as (I) for jaw position.

In the licking task, mice preemptively oriented their jaws and noses toward the lick target on fast RT trials but not on slow RT trials (Figure 4B-C). For some animals, the shape of orofacial features or the degree of separation varied from the others (Figure 4D-E, e.g. mouse LM04 vs LM03), suggesting that there is a degree of individual variability to how mice get ready. Nevertheless, preemptive orienting of jaw and nose toward the lick target was reliably observed across all mice (Figure 4D-E). In contrast, we did not observe reliable differences in the back and tail positions between fast and slow RT trials (Supplemental Figure 3A). In the reaching task, motor readiness was reflected in a subtle displacement of the forelimb toward the reach target (Figure 4G). The forelimb was displaced on fast RT trials but not on slow RT trials. Preemptive forelimb displacement toward the reach target was reliably observed in all mice (Figure 4I). Mice also displaced their jaws and noses on fast RT trials (Figure 4H), but not in a reliable fashion (Figure 4J and Supplemental Figure 3B). Jaws and noses were more protracted relative to the baseline position in some mice but were more retracted in other mice (Figure 4J, e.g. mouse RM02 vs. RM03), and the displacement was not always oriented toward the reach target (Figure 4J, e.g. mouse RM02). Finally, differences in RT were not reliably reflected in the back and tail positions (Supplemental Figure 3B), although some mice did arch their backs in preparation during the reaching task. Altogether, these data show that motor readiness was most reliably reflected in the task-relevant body features (jaw and nose for the licking task and forelimb for the reaching task) compared to other body features.

The displacement in task-relevant body features was evident at the end of the delay epoch, which was well before movement initiation (Supplemental Figure 4A). Importantly, when we analyzed the position of task-relevant body features at movement onset, the slow RT trials still did not show apparent deviation from the baseline position (Supplemental Figure 4B). These analyses show that the preemptive body feature displacement was not related to movement initiation, because movement initiation in absence of motor planning did not require the same body feature displacement on slow RT trials. Therefore, the preemptive body feature displacement on fast RT trials likely reflected the animals’ motor readiness.

Does the body feature displacement track the state of motor readiness in real-time? To investigate this, we examined the probe trials (Figure 5A). On probe trials, the Go cue came at an unexpected time, thereby catching the animals at different states of readiness at the time of its arrival (Figure 2C-D). The average jaw or forelimb position at the Go cue for different probe trial conditions thus revealed the time course of body feature displacement. The jaw and forelimb were chosen because they most reliably reflected motor readiness in their respective tasks. The change in task-relevant body features occurred gradually during the delay epoch: with increasing preparation time, mice showed greater amounts of body feature displacements (Figure 5B-C). To quantify this, we computed a delta-jaw or delta-forelimb value to represent the degree of deviation from the baseline position (Figure 5D-E, Methods). The mean delta values increased as a function of delay duration, peaking at 1.5-1.6 s of preparation time before decreasing slightly with longer delay duration (Figure 5D-E). This time course of body feature displacement mimicked the RT savings as a function of delay duration (Figure 2C-D). Delta-jaw or delta-forelimb values in control trials at the corresponding time points showed a similar time course (Figure 5D-E). Together, these results suggest a progression of readiness over the course of motor planning that could be visualized and quantified in real time using the body feature readout.

**Figure 5:**
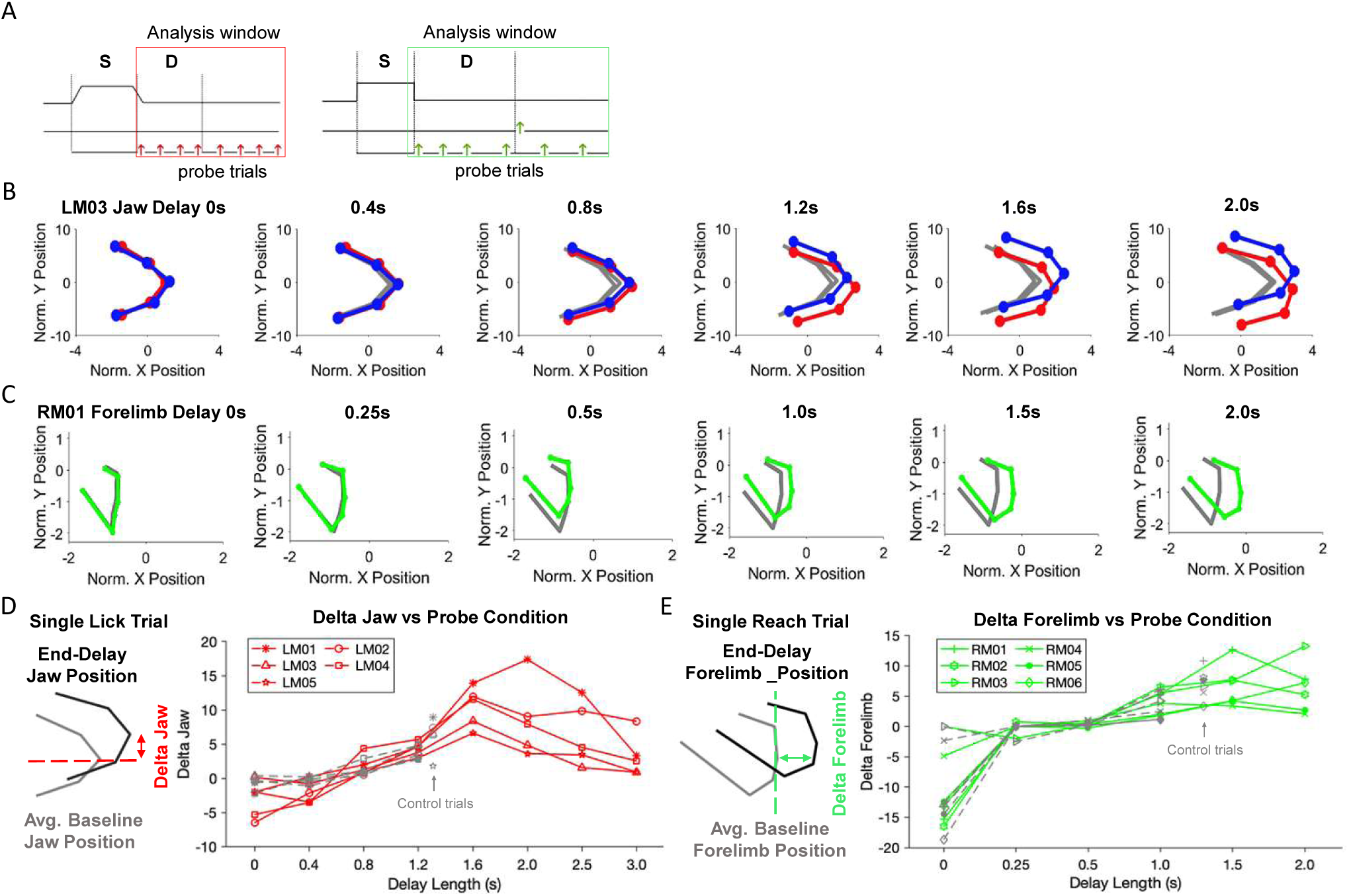
Body feature readout of motor readiness reveals time course of motor planning. (A) Task timeline and analysis window, end of the delay epoch or before the Go cue on probe trials. (B) The position of the jaw in the licking task for one mouse. Plots show the average position of the jaw before the Go cue on probe trials when the Go cue was given at different times of the task. The length of the delay epoch is indicated for each probe trial condition, which is defined as the interval between the pole offset and the Go cue. The mouse shows a progressive orienting of the jaw toward the lick target, which reaches a “ready” position at the expected time of the Go cue (control trial delay duration, 1.3s) and holds at the ready position. (C) Same as (B) for the forelimb position in the reaching task. Data from one mouse. The mouse pre-emptively displaced the forelimb toward the reach target, which reaches a “ready” position at the expected time of the Go cue (control trial delay duration, 1.3s) and holds at the ready position (D) *Left*, quantification of jaw displacement (delta-jaw). A baseline position is computed by averaging the position of the jaw at the start of the delay epoch across all trials (gray). The average deviation in jaw position in the Y dimension is then calculated for each trial. *Right*, delta-jaw values before the Go cue for all probe trial conditions in the licking task. Individual red lines and symbols show individual mice. Gray dashed lines show delta-jaw values on control trials at matched time points in the task (the delay epoch lasted up to 1.3s in control trials). (E) Same as (D) for the forelimb position in the reaching task. Delta-forelimb values are calculated in the same way as delta-jaw values, but in the X dimension.

These results show that motor planning is reflected online in a preemptive displacement of task-relevant body features. The amount of body feature displacement corresponds to different degrees of motor readiness. Interestingly, these data show that mice gradually prepare movements to reach the maximum readiness at the expected time of the Go cue (Figure 5D-E), suggesting that mice learn to anticipate the timing of upcoming movement.

### Body features reflect degree and content of motor readiness on single trials

To further test whether body features reflected the degree of motor readiness online, we asked whether the degree of body feature displacement could predict animals’ RT on single trials. Figure 6B and F show example trials with fast and slow RT in the licking task and reaching task. The fast RT trials were accompanied with a high degree of jaw and forelimb displacement in their respective tasks at the end of the delay epoch, and the slow RT trials had little body feature displacement (Figure 6B and F, delta-jaw or delta-forelimb). To quantify this relationship at the level of single trials, we computed Pearson’s correlations between the delta-jaw or delta-forelimb values and RT across individual trials within each session. Across all trials, delta-jaw and delta-forelimb were anti-correlated with RT (Figure 6C and G). Interestingly, in the licking task, when mice licked to the direction opposite from that instructed by the sensory stimulus, delta-jaw was positively correlated with RT (Figure 6C, error trials; positive correlation corresponds to jaw orienting to the opposite direction from the instructed lick target). This suggests that the body feature readout faithfully reflected the animals’ prepared movements even for erroneous actions.

**Figure 6:**
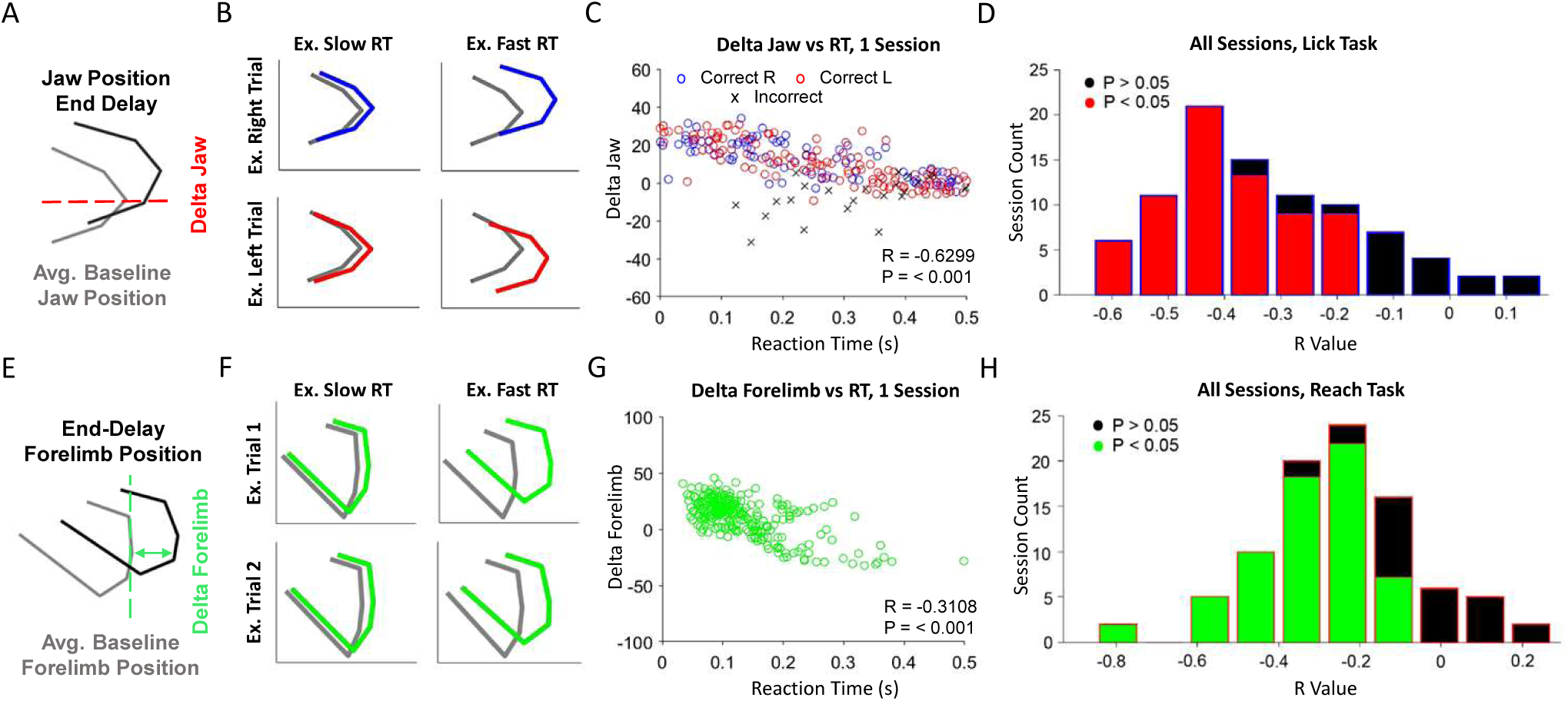
Body feature readout of motor readiness predicts reaction time on single trials. (A) Quantification of jaw displacement (delta-jaw) in the licking task. (B) Example single trial delta-jaw positions for slow and fast RT trials. Lick left trials are shown in red and lick right trials are shown in blue. Gray lines show the baseline position at the start of the delay epoch. (C) Single trial delta-jaw versus reaction time for the licking task. Data from one example session. Red, lick left trials; blue, lick right trials. Circles, correct trials; Crosses, incorrect trial. Because the licking task involved two lick directions, the sign of the delta-jaw values is flipped for lick left trials so that greater deviation from the baseline position always corresponded to greater positive values. Pearson’s correlation and P-values are shown. (D) Histogram of Pearson’s correlation for all sessions, calculated as in (C). Data from sessions and all mice in the licking task. Red shading indicates the sessions with significant correlation (P<0.05). (E) Quantification of forelimb displacement (delta-forelimb) in the reaching task. (F) Example single trial delta-forelimb positions for slow and fast RT trials. (G)Single trial delta-forelimb versus reaction time for the reaching task. Data from one example session. (H) Histogram of Pearson’s correlation for all sessions and all mice in the reaching task. Green shading indicates the sessions with significant correlation.

Across all mice and all sessions, significant anti-correlation between body feature displacement and RT was present in 84.2% of the licking task sessions (64 out of 76) and 77.8% of the reaching task sessions (56 out of 72; Figure 6C and G; mean correlation coefficients, −0.329 ± 0.176 for the licking task; −0.249 ± 0.200 for the reaching task). In these “conventional” sessions, the jaw or forelimb positions thus predicted RT on single trials in their respective tasks (Supplemental Figure 5A-B and I-J). In the licking task, the nose position (delta-nose) also showed an anti-correlation with RT (Supplemental Figure 5C-D), which confirmed that motor readiness was expressed in an orofacial orienting toward the lick target that could be read out in either the jaw or the nose on single trials (Figure 4D-E). Intriguingly, in a small subset of “unconventional” sessions, body feature displacement did not predict RT (Figure 6D and H, Supplemental Figure 5E-H and K-L). In those sessions, mice appeared to be unprepared ubiquitously across all trials regardless of RT (Supplemental Figure 5E and K). We hypothesized that this phenomenon might be tied to the animals’ motivation state. To test this, we compared each session’s correlation coefficient to the amount of water mice consumed in the session, or to the number of trials mice performed in the session (Supplemental Figure 5M and N). These measures of motivation showed that mice displayed a lesser degree of motor readiness in the body feature readout in less motivated sessions. Overall, these data show that the body feature readout reflects the degree of motor readiness and is influenced by motivation state.

In addition to reflecting the degree of motor readiness (i.e., measured in RT), the body feature readout also reflected the content of motor readiness, i.e., what specific action was prepared. This was most clearly illustrated in the licking task, wherein mice prepared different directions of licking on different trials. On probe trials, the Go cue arrived at an unexpected time to catch mice in a particular state of motor readiness. This allowed us to examine what action mice was ready for by observing their movement execution upon the Go cue. For some mice, early arrival of the Go cue resulted in a curious bias in lick direction where mice always licked to one direction regardless of the sensory instruction, resulting in reduced performance for one trial type (Supplemental Figure 6A shows a mouse with lick right bias). This suggests that the mouse adapted a strategy where it was always prepared for one direction of licking by default (“default lick direction”), and it corrected this default lick direction by actively preparing for the other lick direction during the delay epoch. Consistent with this notion, we observed little savings in RT for the default lick direction across probe trial conditions, but substantial RT savings for the other lick direction with preparation time (Supplemental Figure 6B). This strategy of preparing for specific lick direction was reflected in the body feature readout: jaw position did not differentiate fast and slow RT trials for the default lick direction; in contrast, jaw position was selectively displaced for the other lick direction during the delay epoch, reflecting active motor planning for the specific action (Supplemental Figure 6C). Different mice showed default lick direction biases toward different lick targets (Supplemental Figure 6G-I show a mouse with lick left bias). These idiosyncratic biases were more pronounced in the early behavioral sessions but diminished in the late sessions, suggesting that highly trained mice prepared for both lick directions based on sensory instructions (Supplemental Figure 6D-E and J-K). Correspondingly, the preemptive jaw displacement occurred for both lick directions in highly trained mice (Supplemental Figure 6F and L).

Altogether, these results show that body feature readout can reveal both the degree and content of motor readiness. The online readout predicted mice’s RT on single trials and revealed individual variability in each mouse’s motor planning strategy.

### Body feature readout reveals simultaneous motor planning

Given that the body feature readout can reflect motor planning of specific actions online, we next used these measures to examine how mice plan movements in dynamic and ambiguous contexts. Our goal was to create a situation of uncertainty in which mice might benefit from preparing multiple actions simultaneously, which could allow them to rapidly correct a missed movement by switching to the other movement with little RT penalty. Previous studies in humans and monkeys suggest that subjects may prepare multiple potential actions in parallel to rapidly select among them (Cisek and Kalaska, 2005, 2010; Cui and Andersen, 2011; Klaes et al., 2011). However, it remains unresolved whether subjects can simultaneously exhibit motor readiness for multiple actions (Gallivan et al., 2018). Alternatively, subjects may plan one movement at a time (Nashed et al., 2017), even in cases where they must make rapid corrections or make sequences of movements (Ames et al., 2014; Zimnik and Churchland, 2021).

We created a combined licking-reaching task (Figure 7A). Mice were trained to associate an auditory instruction with a specific movement. A 2kHz tone instructed a lick, while a 10kHz tone instructed a reach. Mice initiated each trial by holding a resting bar for 0.5s. At the beginning of each trial, a spout moved into one of two positions: a position within reach of the tongue (“lick trial”) or a position where the tongue could not reach (“reach trial”). After this, the instruction tone played for 0.5s. After a delay epoch (1.3s), an auditory Go cue indicated the start of the response epoch (pure tone, 3.7kHz, 0.1s duration) in which mice could either lick or reach for the spout to obtain a drop of water reward (Figure 7A). If mice missed the target upon the first attempt, they could correct the action by making the other movement to obtain the reward. To examine motor planning under uncertainty, we used an uninstructive tone (Figure 7B, a mashup of 2kHz and 10kHz tones, Methods) and we removed all cues that were informative about the spout location. Mice performed the task in the dark and the whiskers were trimmed. The task thus probed preparation of either licking or reaching and subsequent movement executions.

**Figure 7:**
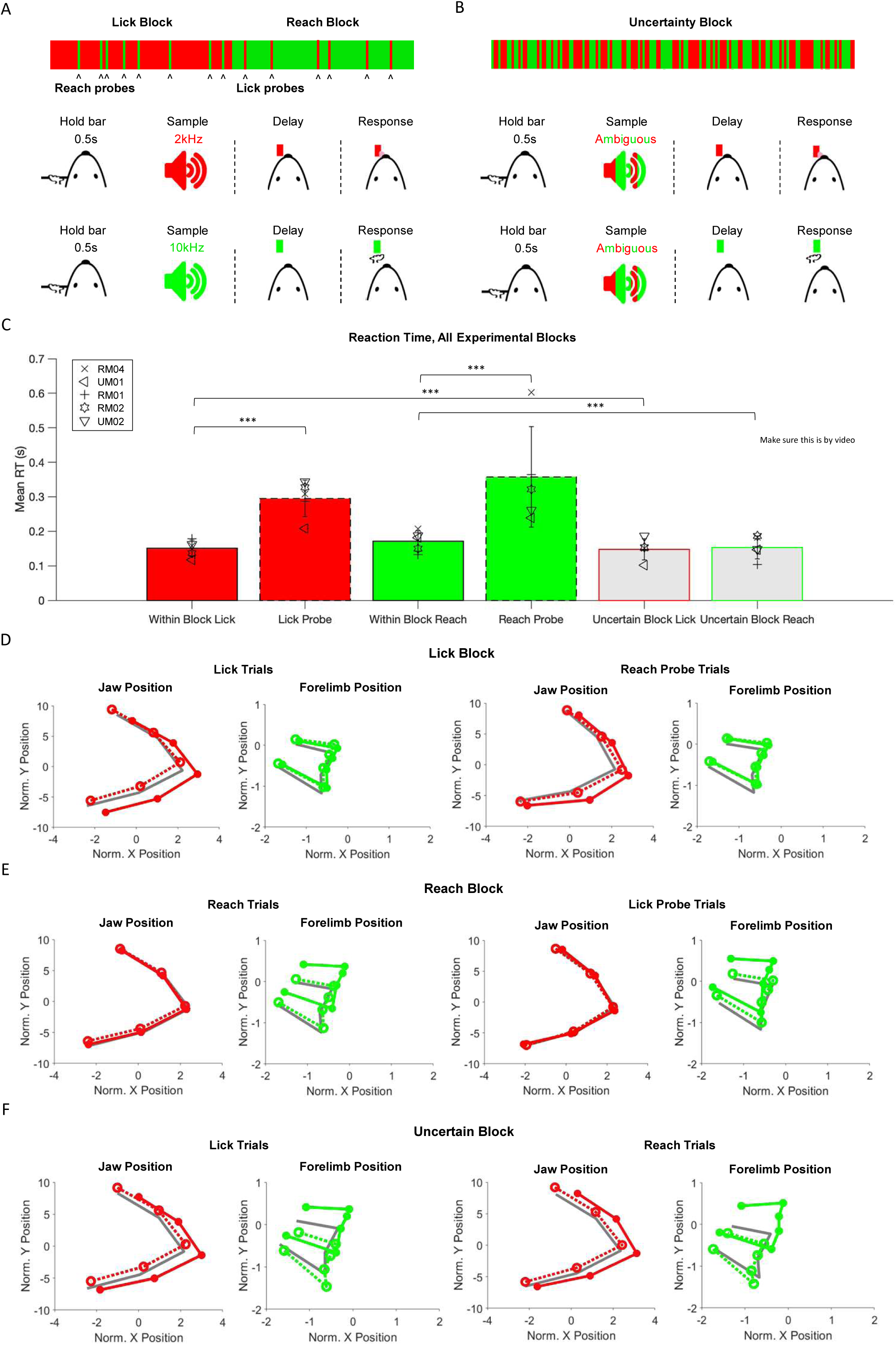
Body feature readout of motor readiness reveals simultaneous motor planning of multiple actions. (A) Combined licking-reaching task, single movement blocks. *Top*, block structure. In lick blocks, most trials are lick trials (red) with a few probe reach trials (green, 4% of total). In reach blocks, most trials are reach trials with a few probe lick trials (4% of total). *Bottom*, trial structure. Mice hold a resting bar for 0.5s to begin each trial. A single spout moves into a position reachable by the tongue (lick trials) or forelimb (reach trials). A low-frequency tone (2kHz, 0.5s) instructs licking or a high-frequency tone (10kHz, 0.5s) instructs reaching. After a 1.3s delay, the Go cue plays (3.7kHz, 100 ms) and mice can lick or reach for water. (B) Combined licking-reaching task, uncertain block. Every trial has a 50% probability of being a lick or reach trial, and the movement instructing auditory tone is an ambiguous mashup of the licking and reaching cues. (C) Reaction time (RT) for all conditions. Bars, mean; error bars, SD across mice; symbols, individual mice. In the single-movement blocks, RT is fast for the expected action but slow for the unexpected action on probe trials. In the uncertain blocks, RT is fast for both actions. (D) Average jaw and forelimb positions at the end of the delay epoch in the lick block. *Left*, lick trials within the lick block. *Right*, probe reach trials within the lick block. Solid colored lines, average positions on fast RT trials. Dashed lines, average positions on slow RT trials. Gray, average positions at the start of the delay epoch (baseline). Within the lick block, mice show body feature of motor readiness for licking on both lick trials and probe reach trials. Thus mice ignore the movement-instructing tone and do not prepare for reaching. (E) Average jaw and forelimb positions in the reach block. Within the reach block, mice show body feature of motor readiness for reaching on both reach trials and probe lick trials. Thus mice prepare for reaching and do not prepare for licking, (F) Average jaw and forelimb positions in the uncertain block. *Left*, trials in which the spout is in position for licking (“lick trials”). *Right*, trials in which the spout is in position for reaching (“reach trials”). Mice show behavioral feature of motor readiness for both licking and reaching regardless of trial type.

As a baseline, we first measured motor planning of single movements without uncertainty (Figure 7A). In a block of trials, mice were nearly always instructed to lick (“lick block”) or reach (“reach block”). Occasionally (on 4% of the trials), we tested probe trials in which the mice were instructed to make the other movement (Figure 7A). Within a block, mice always prepared for the expected movement and failed to prepare for the unexpected movement in probe trials. RT was faster for the expected movement within block compared to the same movement when instructed unexpectedly in probe trials (Figure 7C), and the resulting movement was more accurate on fast RT trials (Supplemental Figure 7A). Motor planning was reflected in the task-relevant body features. In the lick block, mice preemptively oriented their jaws toward the target, but did not move the forelimb (Figure 7D). In the reach block, mice preemptively displaced their forelimb toward the target, but did not orient their jaws (Figure 7E). On probe trials, the body feature readout indicated that mice prepared for the wrong movement type (Figure 7D-E), thus explaining the increased RT on probe trials (Figure 7C). These results confirmed that task-relevant body features could be used to readout motor planning of specific actions (i.e., licking versus reaching).

After mice learned to perform the task with a high degree of accuracy, they entered into an ambiguous block where movement instructing cues were removed and every trial had a 50% probability of being a lick trial or a reach trial (Figure 7B). In this uncertain context, the body feature readout revealed that mice simultaneously prepared for both licking and reaching. On fast RT trials, mice oriented their jaws while also displacing their forelimbs toward the target location (Figure 7F). Mice preemptively displaced both body features regardless of which action they ultimately performed (Figure 7F, lick trials vs. reach trials), and both body features were simultaneously displaced on single trials (Supplemental Figure 7B-C). However, simultaneous motor planning was not ubiquitous across all mice: 3 out of 5 mice tested showed such signatures of simultaneous motor planning while the other 2 mice adapted a strategy of “defaulting” to always preparing one movement type (Supplemental Figure 7B). To validate mice indeed prepared for both movement types, we examined their RT. RT was defined by whichever movement mice initiated first after the Go cue (licking or reaching). Mice exhibited fast RT for both licking and reaching in the uncertain block (Figure 7C), indicating that they were simultaneously ready for both actions. At the level of individual trials, the degree of body feature displacement predicted RT of both licking and reaching regardless of which action the mice took (Supplemental Figure 7C).

Thus, body feature readout revealed that mice could exhibit motor readiness for multiple actions in uncertain situations, which enabled rapid selection of prepared actions.

## Discussion

We combined videography and delayed response task paradigms in an autonomous behavioral system to measure motor planning in non-body-restrained mice. We observed reaction time (RT) savings with preparation time and the resulting movements were more precise (Figures 2-3). Motor readiness was reflected online in task-relevant body features as mice prepared for specific movements (Figure 4). Both the specific action prepared and the degree of motor readiness could be reliably read out from body features well before movement initiation, even on single trials (Figure 6). This online readout of motor readiness revealed that mice anticipated the timing of the Go cue: they gradually prepared movement to reach maximum readiness at the expected time of movement initiation (Figure 5). Body features further revealed behavioral evidence of simultaneous motor planning of multiple actions under uncertain situations (Figure 7).

Our study makes 3 contributions. First, we provide a detailed behavioral characterization of motor planning in mice, an increasingly used model to study motor planning. Preparatory activity has been reported in mouse motor cortex and in connected brain regions (Svoboda and Li, 2018). Experiments using mice have largely focused on the contributions of individual brain regions to preparatory activity (Guo et al., 2014a; Li et al., 2015; Guo et al., 2017; Hasegawa et al., 2017; Gao et al., 2018; Chabrol et al., 2019; Wang et al., 2021), but behavioral measurements of motor planning have thus far been lacking. RT saving has long been used as a metric of motor planning along with movement accuracy or performance in humans and monkeys (Rosenbaum, 1980; Riehle and Requin, 1989; Churchland et al., 2006a; Churchland et al., 2006b; Michaels et al., 2015; Dahan et al., 2019). We find that motor planning in mice produces the same beneficial effects on subsequent movements by these classical measures, thus validating the mouse as a model to study neural mechanisms of motor planning.

Second, we introduce and validate the body feature readout as a new way to measure motor readiness in real time. Classic measures of motor planning such as RT, accuracy, or performance are obtained from movement execution. They therefore cannot reveal the degree of motor readiness before movement onset, and they provide little information about what actions are prepared (the content of motor readiness). A recent study in mice found that motor planning and behavioral state can be reflected in whisking patterns (Dominiak et al., 2019). Here we show that task-relevant body features can be used to read out both the degree of motor readiness and the specific actions prepared in real time, which reveals the time course of motor planning as a gradual process that anticipates the expected timing of movement initiation (Figure 5).

Third, body feature readout reveals that mice are capable of preparing for two actions simultaneously when the movement goal is uncertain. Motor planning has been largely studied as a serial process that prepares one movement at a time, leading to the optimal initial condition hypothesis that poises unique readiness activity states for specific actions (Afshar et al., 2011; Shenoy et al., 2013; Gallego et al., 2017). Little is known about how motor planning extrapolates to dynamic and uncertain contexts that often require rapid changes in movement execution (but see (Cisek and Kalaska, 2005; Klaes et al., 2011; Ames et al., 2014; Zimnik and Churchland, 2021)). Simultaneous motor planning of multiple potential actions may allow rapid selections of prepared actions with little RT penalty (Cisek and Kalaska, 2010; Gallivan et al., 2018). Uncovering behavioral evidence of simultaneous motor planning in mice enables analysis of its neural substrates.

### Relation to previous studies

Recent studies show that ongoing movement is prominently represented in brain-wide neural activity (Musall et al., 2019; Stringer et al., 2019; Salkoff et al., 2020; Zagha et al., 2022), including in brain regions that participate in motor planning (Musall et al., 2019). Our data suggest that ongoing movements and posture changes may be an inseparable part of motor planning: motor readiness is strongly correlated with the degree of preemptive body feature changes. This raises intriguing questions about the role of these preemptive posture changes in motor planning. The preemptive posture changes may simply reflect central processes related to motor planning without directly contributing to future movements. Alternatively, motor planning may be mediated by reciprocally coupled sensorimotor loops that span the brain and the body, whereby reafferent activity caused by the posture changes in turn contributes to ongoing motor planning. Definitively resolving which specific activity contributes to motor planning will require causal manipulations.

In the delayed response tasks with a fixed delay duration, we find that mice anticipate the future timing of the Go cue: mice gradually prepare movement to reach maximum readiness at roughly the expected time of the Go cue, after which the RT saving effect remains or even slightly diminishes (Figure 2C-D and 5D-E). This optimal time window of maximum motor readiness varies slightly across individual mice. Similar optimal motor planning time windows have been previously reported in humans (Dahan et al., 2019) and monkeys (Michaels et al., 2015).

Considering this temporal feature of motor planning within the optimal initial condition framework (Shenoy et al., 2013), there seem to be two separate processes involved. Motor planning of specific movements is thought to be associated with different patterns of neuronal activity in motor cortex (Tanji and Evarts, 1976; Riehle and Requin, 1989; Alexander and Crutcher, 1990; Crammond and Kalaska, 2000; Cisek and Kalaska, 2005; Churchland et al., 2010). A preparation process brings activity into a specific region in activity space, i.e., an activity state that constitutes a fast-RT initial condition for specific movement (Afshar et al., 2011; Shenoy et al., 2013). This process may specify the identity of the actions prepared. In addition, previous studies also revealed a ramping activity in motor cortex and connected brain regions that increases to a fixed threshold before movement onset (Hanes and Schall, 1996; Roitman and Shadlen, 2002; Maimon and Assad, 2006; Tanaka, 2007; Murakami et al., 2014; Thura and Cisek, 2014). Furthermore, the slope of the ramping can flexibly adapt to the task timing (Wang et al., 2018; Inagaki et al., 2019). Ramping activity is prominent in mouse ALM during motor planning (Inagaki et al., 2019; Finkelstein et al., 2021) and the ramp amplitude correlates with RT (Yang et al., 2022). The ramping activity may reflect a separate process that determines the expected timing of movement initiation or hazard rate for the upcoming Go cue (Janssen and Shadlen, 2005; Cisek et al., 2009; Remington et al., 2018).

Several lines of evidence support this description of motor planning. In delayed response tasks where the timing of the Go cue is fixed and predictable, preparatory activity reaches specific readiness activity states via a “prep-and-hold” process reflected in the slow ramping reaching a threshold. However, reaching this activity state is not sufficient to trigger movement initiation autonomously. If there are probe trials in which the Go cue unexpectedly arrives later, the activity holds at the threshold in absence of the Go cue (Tanaka, 2007). The temporal profile of motor readiness measured in our study also supports this notion, where mice reach maximum readiness at the expected time of the Go cue but continue to exhibit RT savings in probe trials with longer than expected delay (Figure 2C-D and 5D-E). In delayed response tasks where the timing of the Go cue is unpredictable (i.e., no timing to be learned). The activity ramps to the threshold quickly and holds at the threshold (Ames et al., 2014; Peixoto et al., 2018; Inagaki et al., 2019). Activity driving movement (output-potent activity) seems to be triggered by a separate internal Go signal (Inagaki et al., 2022), at least in tasks that involve cue-triggered movement.

The initial condition in activity space may therefore represent a ‘readiness buffer’ that holds the prepared action without initiating it and can be updated if needed (Cisek and Kalaska, 2010). For example, if a different instruction is given after activity reaches an initial condition, the activity can hop over to a different initial condition in activity space (Bracewell et al., 1996; Ames et al., 2014), presumably reflecting a change of motor plan. In this framework, since unique locations in activity space correspond to readiness for specific future actions, the finding that mice can simultaneously plan multiple actions open intriguing questions about how such activity state could be supported by preparatory activity (Figure 7) (Meirhaeghe et al., 2022). Future studies using population neural recordings during simultaneous motor planning is needed to shed light on this question. Targets for such recordings may include the rostral forelimb area (RFA) and ALM, which exhibit extensive spatial overlap with each other (Svoboda and Li, 2018). ALM is involved in motor planning of directional licking (Guo et al., 2014a; Li et al., 2015; Bollu et al., 2021; Xu et al., 2022) whereas RFA is thought to be involved in motor planning of forelimb movement (Hasegawa et al., 2017; Morandell and Huber, 2017).

A key function of the motor system is to navigate environmental dynamics and uncertainties (Cisek and Kalaska, 2010; Wolpert and Landy, 2012; Cisek and Pastor-Bernier, 2014; Dunning et al., 2015; Merel et al., 2019). Cisek and colleagues postulate that motor planning of potential actions are continuous, parallel processes that not only lead up to movement initiation but also ongoing during movement execution (Cisek and Kalaska, 2010). Previous studies in humans and monkeys have examined motor planning under uncertain situations and whether subjects plan multiple potential actions in parallel (Cisek and Kalaska, 2005; Cui and Andersen, 2011; Klaes et al., 2011; Gallivan et al., 2016; Nashed et al., 2017; Gallivan et al., 2018). Our finding that mice can simultaneously exhibit motor readiness for multiple actions adds to this line of work. The ability to simultaneously prepare multiple actions and continuously adjust motor plans over ongoing movements may allow animals to navigate dynamic environments (Bollu et al., 2021; Xu et al., 2022). It will be of interest to determine how such dynamic motor planning processes are mediated by neural circuit dynamics.

## Acknowledgements

We thank Javier Medina, Matthew McGinley, Jeff Yau, Jacob Reimer for insightful discussions and Takashi Sato, Michael Economo, Jae-Hyun Kim, Weiguo Yang, Jaclyn Birnbaum, Munib Hasnain, Runbo Gao for comments on the manuscript. This work was funded by the Robert and Janice McNair Foundation, Whitehall Foundation, Alfred P. Sloan Foundation, Searle Scholars Program, Pew Scholars Program, NIH NS104781, NS112312, NS113110, McKnight Foundation, and Simons Collaboration on the Global Brain.

## Author contributions

EM and NL conceived and designed the experiments. EM performed the experiments. JL developed the hardware for behavioral experiments. JC developed the video acquisition and analysis pipeline. EM analyzed the data. EM and NL wrote the paper.

## Materials and methods

### Animals

Subjects (n = 13) consisted of male and female wildtype C57BL6/J mice aged 2-7 months, obtained from Jackson Laboratory. 5 mice were tested in the delayed response licking task. 6 mice were tested in the delayed response reaching task. 5 mice were tested in the combined licking-reaching task, including 3 mice that were previously tested in the delayed response reaching task. All experiments and procedures were carried out in accordance with protocols approved by the Institutional Animal Care and Use Committees at Baylor College of Medicine (protocol AN-7012). Mice were singly housed at constant temperature (22 ± 1 °C) and humidity (30-55%) under a 12:12 reversed light/dark cycle. Mice were tested during the dark phase and all experiments were carried out in darkness. Before behavioral training began, mice were water-restricted for 5-7 days. During restriction, mice were handled daily and received ∼1mL of water per day until they reached a stable body weight. On experimental collection days, mice were tested in experimental sessions lasting 2 hours each where they received water via engagement in behavioral tasks. On days not tested, mice received 0.6-1mL of water. Mice were given supplementary water if they did not maintain a stable body weight (Guo et al., 2014b). Mice voluntarily engaged in the delayed response tasks in an autonomous behavioral system (Hao et al., 2021). Mice that became too old for testing or failed to engage in the task for a prolonged period were removed from the study (i.e., not included in the mouse count above; 1 mouse in the licking task, 2 mice in the reaching task, and 2 mice in the combined licking-reaching task).

### Headbar implant surgery

All surgical procedures were carried out aseptically under 1-2% isoflurane anesthesia. Buprenorphine Sustained Release (1 mg/kg-1) was administered two hours before surgery. Additionally, mice were given a mixture of bupivacaine and lidocaine topically before incision. Meloxicam Sustained Release (4 mg/kg-1) was used for postoperative analgesia. After surgery, mice were given at least 3 days to recover with *ad libitum* access to water before water restriction began.

Mice were fitted with a custom headbar to allow for voluntary head-fixation within the behavior apparatus (see (Hao et al., 2021)). Briefly, the scalp and periosteum over the dorsal surface of the skull were removed and the position of Bregma and Lambda were noted. The skull was gently scraped using a dental drill to facilitate headbar attachment. The anterior edge of the headbar was aligned with Lambda (approximately over the cerebellum) and attached to the skull with layer of cyanoacrylate adhesive (Krazy Glue, Elmer’s Products). A thin layer of clear dental cement (1223CLR, Lang Dental) was applied over the adhesive, covering the entire exposed skull.

### Behavior apparatus

Mice were tested in an autonomous behavioral system (Fig 1E-G). The details of the autonomous behavioral system are described in (Hao et al., 2021). Briefly, the system consisted of a behavioral test chamber attached to an empty home-cage. Behavioral apparatuses for the delayed response task were inside the test chamber. A 25×25 mm opening (“headport”) connected the home-cage to the test chamber. Mice engaged in the delayed response task by accessing the test chamber through the headport (Fig 1F-I). Two snap action switches were mounted on both sides of the headport. Mice were fitted with a custom headbar. Upon head entry, the headbar triggered the switches and two pneumatically driven pistons clamped the headbar for head-fixation. In daily sessions, mice were place into the behavioral system for 2 hours in which they self-engaged in head-fixation and delayed response tasks for water rewards.

In the licking task, a motorized lickport with two spouts (5mm apart) was placed in front of the headport. In the reaching task, only a single spout was presented. Two orthogonal linear motors (L12-50-100-12-I and L12-30-50-12-I, Actuonix) controlled forward/backward and left/right movement of the spouts. Each spout was electrically coupled to a custom circuit board that detected contact via electrical circuit completion (Slotnick, 2009; Guo et al., 2014b). Water rewards were dispensed by two solenoid valves (LHDA1233215H, Lee Co.).

The sensory stimulus for the licking task was a mechanical pole (1.5 mm diameter) on the right side of the headport. The pole was motorized by a linear motor (L12-30-50-12-I, Actuonix) and presented at different location to stimulate the whiskers. The motorized pole was attached to an air piston (6498K999, McMaster), driven by a 3/2-way solenoid valve (196847, Festo), which moved the pole vertically into the reach of the whiskers. The entire pole mechanism was mounted on a holder attached to the behavioral test chamber. The auditory Go cue was provided by a piezo buzzer (3.7 kHz, cpe163, Mouser).

In the reaching task, a hole (10mm) was drilled into the left side of the headport to allow the left forelimb to fit through (Fig 1G). A horizontal resting bar was placed along the bottom edge of this hole, roughly 20-25mm below the mouse’s headbar. The resting bar was electrically coupled to the circuit board to detect touch (similar to the lickspout above), which was monitored by an internal timer in the autonomous behavioral system’s Arduino board. Mice rested their forepaws on the resting bar when not reaching. The sensory stimulus for the reaching task was an auditory pure tone (10kHz, 0.5s). The auditory Go cue was the same as the licking task (3.7kHz, 0.1s).

Training parameters and behavioral data were stored on a SD card which could be inserted into the Arduino board that controlled the behavioral system. These data included detailed information for each trial such as trial number, trial type, trial outcome, number of licking or reaching events, and events related to head-fixation. Important aspects of behavior data such as the timing of licking or reaching events were timestamped using a real time clock (RTC). Behavior was monitored in real time using a custom written MATLAB GUI (MathWorks) (Hao et al., 2021).

### Video acquisition

Three CMOS cameras (CM3-U3-13Y3M-CS, Point Grey) were used to track orofacial and body movements of the mouse from the side, bottom, and front views (Fig 1H-I). The front camera was positioned over the headport (field of view size: 352 × 256 pixels, width × height). The bottom camera was situated under the lickport at the bottom of test chamber (384 × 288 pixels). The side camera was positioned to the side of the home-cage (480 × 256 pixels). Mice performed the task in complete darkness, and videos were acquired under infared LED illumination (940nm, SM-01-R9, LuxeonStar) mounted on heat sinks (N30-25B, LuxeonStar). Videos were recorded at 300 frames/s (side camera) or 350-360 frames/s (front and bottom cameras). Video acquisition started upon the start of each trial and stopped at the end of each trial. Videos therefore varied in length depending on the length of the trial. A custom written software controlled the video acquisition.

### Behavioral tasks and training

#### Licking task

The delayed response licking task and training procedures have been described previously (Guo et al., 2014b; Hao et al., 2021). The stimulus was a 3D-printed vertical pole (1.5mm diameter) presented at either an anterior or posterior location (Figure 2A). The two possible positions were spaced 5mm apart along the anterior–posterior axis, with the posterior pole position 5mm from the whisker pad. At the beginning of each trial, the pole moved into reach of the whiskers (travel time: 0.2s) and remained for 1s before retracting (retraction time: 0.2s). The sample epoch was thus defined as the time between pole movement onset to 0.1s after the pole retraction onset (sample epoch: 1.3s). Mice discriminated the pole location with their whiskers, typically touching the pole at different locations using different whiskers. The anterior pole position instructed a left lick while the posterior position instructed a right lick. The delay epoch (1.3s for control trials) followed and was ended by an auditory Go cue (pure tone, 3.7kHz, 0.1s).

Mice were trained with the fixed 1.3s delay until they reached 75% correct performance, after which probe trials were added. On probe trials, which comprised 10% of total trials, the Go cue arrived randomly at one of 8 pre-defined times relative to the expected time (Fig 2A; possible probe trial delay durations: 0, 0.4, 0.8, 1.2, 1.6, 2.0, 2.5, 3 s). Mice received a water reward (3uL) upon licking the correct lickspout after the Go cue. Incorrect licks triggered a timeout (4s). Licking early during the sample or delay epochs resulted in a brief pause (0.1s). After the pause, the program resumed the trial from the beginning of the epoch in which the mouse licked early. Mice could still obtain water rewards by correctly completing the trial after the early lick. Each trial was separated by an inter-trial interval (2.5s). Trials in which mice did not lick within a 1.5s response window were deemed “ignore” trials and were excluded from analysis. Ignore trials were rare, though some mice exhibited occasional pauses on the shortest-delay probe trials.

#### Reaching task

The reaching task was adapted from a water reaching task developed by (Galinanes et al., 2018). Mice were first screened for handedness using the chamber method described in (Galinanes et al., 2018). Mice were water-restricted and placed into an empty home-cage with a small slit at face-height. The slit was wide enough to fit the forelimb through, but not the snout. A water-dropper was positioned outside the chamber to encourage mice to reach. Mice quickly learned to reach for the water within one 30 min training session. Only left-handed mice were selected for the reaching task because the resting bar and spout were situated on the left-hand side of the autonomous behavioral system (Fig 1G). To acclimate mice to head-fixed reaching, we restrained their headbars in a custom caddy. Their bodies rested in a comfortable position in a restraint tube and their forelimbs were able to freely reach forward. Mice were similarly trained to reach for water using the dropper. Most mice learned this behavior within 2-4 days of training.

Once trained for head-fixed water reaching, mice were placed into the autonomous behavioral system in which they learned to engage in voluntary head-fixation using previously described training protocols (Hao et al., 2021). After learning head-fixation, mice began a training protocol for the reaching task. The spout was positioned to the left of the mouse within reach of the forelimb, but not the tongue. Upon head-fixation, a start cue played (pure tone, 10kHz, 0.5s) followed simultaneously by the Go cue and delivery of a drop of water. Mice typically reached for the water droplet on the spout. After mice consumed the water, a new trial began after an inter-trial interval (2.5s). After mice reliably reached for the water droplet on a majority of trials, a delay epoch was added between the start cue and the Go cue. Initially, the delay epoch was brief (0.3s). The duration of the delay epoch increased by 0.2s as mice progressed through training. Concurrently, mice were trained to initiate reaching from a fixed initial position by placing their forelimb on the resting bar. Initially, mice only needed to touch the bar for a very brief period (0.05s) to initiate a new trial. The holding time requirement increased in intervals of 0.5 to 0.5s at the final stage of training. Mice were allowed to release the resting bar after the trial began.

In the final task, mice learned to hold the bar for 0.5s to initiate each trial and reliably withheld their reaches for the full delay duration of 1.3s until the Go cue. Probe trials were added to compose 10% of the total trials. On these trials, the Go cue came at an unexpected time (Fig 2E). A slightly smaller selection of probe conditions was used based on the results from the licking task (possible probe trial delay durations: 0, 0.25, 0.5, 1.0, 1.5, 2 s). Because the reaching task contained no direction contingency, trials could only be classified as correct or ignore trials. Response time window was limited to 1.5s.

#### Combined licking-reaching task

The combined licking-reaching task consisted of two distinct phases. In phase 1, mice were tested in single-movement blocks in which they were instructed to make a pre-defined movement (Fig 7A). Mice were trained to reach for water using either their tongue or forelimb. Mice initiated each trial by holding the resting bar for 0.5s but were al lowed to release the resting bar after the trial began. On each trial, the spout moved from an unreachable position to a target position. Optimal lick and reach spout positions were determined for each mouse such that the animal could comfortably reach the spout by tongue on lick trials and by extending the forelimb on reach trials. The spout motion duration was matched for the lick and reach trials so that the sound of the spout motion was uninformative about trial type. Once the spout was in position, an instruction cue played. Lick trials were cued by a 2kHz pure tone (0.5s) and reach trials were instructed by a 10kHz pure tone (0.5s). After a 1.3s delay epoch, a water droplet was delivered on the spout simultaneously with the Go cue. There was no penalty for making an incorrect movement (e.g. reaching for water within the lick block). While reaching for water on lick trials was physically possible, mice typically initially missed the spout on their reach trajectory due to spout positioning. Because mice were first trained to reach for water, they initially used their forepaws. To encourage mice to lick rather than using their forelimbs on lick trials, the spout was initially placed directly in front of the mouth in a difficult-to-reach position during training. Once mice learned the association, the spout position was adjusted to slightly below the mouth. Whisker trimming was performed every four days to prevent mice from using whisks to detect the spout location instead of learning the sound cue association. The spout returned to the unreachable baseline position after the trial concluded.

To further ensure that mice prepared the instructed movement, the lick trials and reach trials were provided in blocks of the same trial type (Fig 7A). Once mice reliably licked within lick blocks and reached within reach blocks, data collection began. To examine whether mice prepared for only single actions, probe trials were tested in which the other action were unexpectedly instructed on 4% of the trials (Fig 7A; reaching within lick block or licking within reach block). Mice alternated daily or every other day between lick blocks and reach blocks.

After several days, mice entered phase 2: the uncertain block (Fig 7B). In the uncertain block, every trial had a 50% chance of being a lick or a reach trial, and the start cue was an ambiguous rapid sequence of the two tones (sequence: high-low-high-low in frequency, 0.125s each). Thus, mice had no cues to determine which action they were supposed to prepare.

### Data analysis

#### Video analysis

We used DeepLabCut, an open-source toolbox that allows markerless tracking using a deep neural network, to track manually-defined body parts of interest (Mathis et al., 2018; Nath et al., 2019). Separate models were used for each task-relevant body part (forelimb, tongue, jaw, nose, back, and tail). The development dataset for model training and validation contained manually labeled videos from multiple mice and multiple sessions. 1000-5000 frames were manually labeled within the development sets depending on the body feature, with the tongue tracking model requiring the most labeling. Frames for labeling were automatically selected by the program using k-means clustering to capture a variety of body postures at different timepoints within trials. The labeled frames of the development set were split randomly into a training set and a test set (95% training, 5% test dataset). Training was performed using the default settings of DeepLabCut. All models were trained up to 500,000 iterations with a batch size of one. The trained models tracked the body features in the test data with an average tracking error of less than 2.5 pixels.

#### Reaction time calculation

All reaction times (RTs) were obtained from video analysis. The RT was defined as the time difference between the first frame in which effector motion was detected and the Go cue. The onset of the effector motion was defined differently for lick and reach trials. On lick trials, the first frame in which the tongue appeared after the Go cue was used. On reach trials, we first defined baseline forelimb positions within a time window around the Go cue (approximately 100 ms before and after the Go cue). This baseline time window was selected individually for each mouse depending on their reaction time. We then calculated the mean forelimb position in the X dimension within this time window. The first frame in which the forelimb position in the X dimension reached more than 3 standard deviations above the mean was denoted as a reach onset.

#### Trial sorting by RT

Within each session, the trials were sorted by RT from fastest to slowest. For analysis of movement accuracy (Fig 3) and body feature of motor readiness (Fig 4), the top 33.3% percentile and the bottom 33.3% percentile of the trials were obtained. These were defined as the “fast RT” and “slow RT” trials respectively for all relevant analyses. For probe trial analysis, all trials were pooled together for each condition without dividing into fast and slow RT due to limited number of probe trials.

#### Analysis of body features

For analysis of body features in Figures 4-7 and related supplemental figures, X-Y position plots were generated by averaging the pixel position of the tracked body features across the last five video frames before the Go cue. To account for minute differences in camera position from session to session, position data were normalized in the X and Y dimensions before averaging data across sessions. To normalize, mean marker positions in the X and Y dimensions and their standard deviations were calculated at the session level. This mean was then subtracted from the marker position in the X and Y dimensions on every trial and the result was divided by the standard deviation.

#### Delta-forelimb and delta-jaw computations

As body feature readouts of motor readiness (Fig 5 and 6), delta-jaw and delta-forelimb quantified the deviation of jaw and forelimb from their baseline positions. For each body feature, we determined its centroid position by averaging all the feature markers. The baseline position was calculated by averaging the centroid positions of the first five frames of the delay epoch and averaging across all trials in the session. Delta-jaw and delta-forelimb and were calculated by subtracting the centroid position at the end of the delay epoch (averaging the last 5 frames before the Go cue) from the baseline position. For delta-jaw, we quantified the deviation in the Y dimension to account for lick direction contingency.

For delta-forelimb, we calculated the deviation in the X dimension because the forelimb primarily moved forward in X. Delta-jaw or delta-forelimb values were calculated in individual trials.

### Statistics and reproducibility

No statistical methods were used in advance to determine sample size. All data analysis was performed using MATLAB (MathWorks). All mice were trained and recorded under the same conditions, so randomization of subjects was not necessary. Statistical tests were done using a repeated measures ANOVA test for comparison between more than two groups (MATLAB ‘ranova’) and a Mann-Whitney U-Test for paired data (MATLAB ‘ranksum’). Pearson’s correlations were computed using the MATLAB ‘corr’ function.

---

**Supplemental Figure 1:**
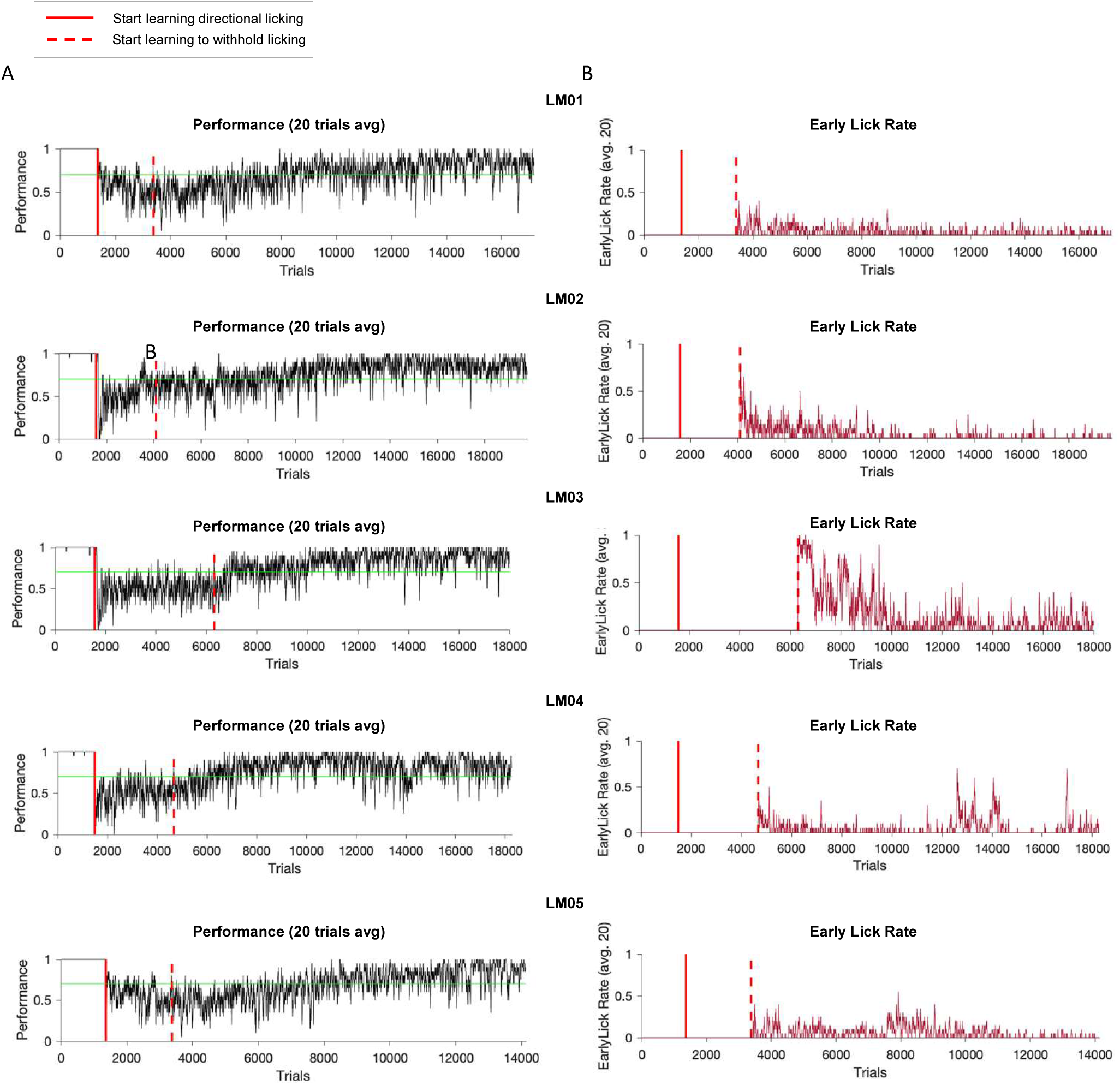
Behavioral performance in the autonomous behavioral system. (A) Behavioral performance across time in the licking task for each of the 5 mice. Red solid lines indicate the start of direction contingency learning (sample protocol). (B) Early lick rates across time for each of the 5 mice. Red dashed lines indicate the start of the delay protocol, in which mice must withhold early licking to avoid receiving a time-out.

**Supplemental Figure 2:**
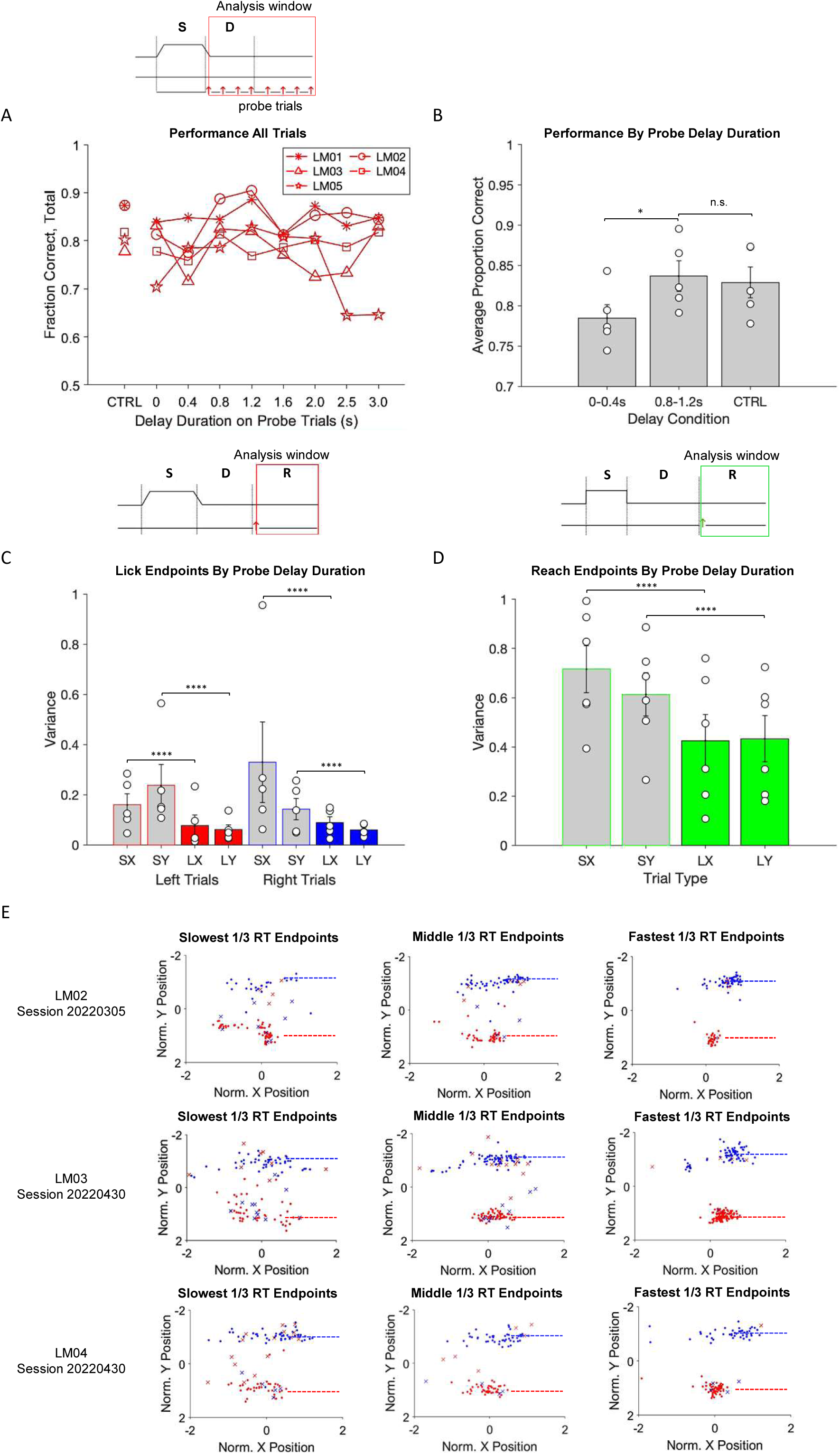
Movement accuracy is correlated with reaction time. (A) *Top*: analysis window, the response epoch (including probe trials). *Bottom*: performance (fraction of correct trials) in the licking task for control trials and for each probe trial condition. Individual lines and symbols show the mean across trials for individual mice. (B) Performance comparison across control, short delay, and long delay conditions. Average performance for each probe category was calculated by combining trials across two representative probe delay conditions (short: 0s-0.4s delays; long: 0.8-1.2s delays). Symbols, individual mice; bars, averages; error bars, SD across mice. (C) *Top*: analysis window, the response epoch, licking task. Bottom: variance of the lick endpoints on short-delay probe trials (S, 0s-0.4s delays) and long-delay probe trials (L, 1.2s-1.6s delays). Variance is calculated along the X and Y directions for each lick direction (lick left and lick right). Symbols, individual mice; bars, averages; error bars, SD across mice. Lick left trials, X variance, short-delay vs long-delay trials: p < 0.00001; Lick left trials, Y variance, short-delay vs long-delay trials: p < 0.00001; Lick right trials, X variance, short-delay vs long-delay trials: p < 0.00001; Lick right trials, Y variance, short-delay vs long-delay trials: p < 0.00001. (D) Same as (C) for the reaching task. Short-delay probe trials are taken as 0s and 0.25s delays while long-delay probe trials are taken as 1.0 and 1.5s delays. Reach endpoints, X variance, short-delay vs long-delay trials: p < 0.00001; Reach endpoints, Y variance, short-delay vs long-delay trials: p < 0.00001. (E) Endpoint distribution for the bottom 1/3, intermediate 1/3, and top 1/3 of the trials ranked by reaction time. Example sessions from mice LM02, LM03, and LM04. Red dots, correct lick left trials. Blue crosses, incorrect lick right trials. Red crosses, incorrect lick left trials.

**Supplemental Figure 3:**
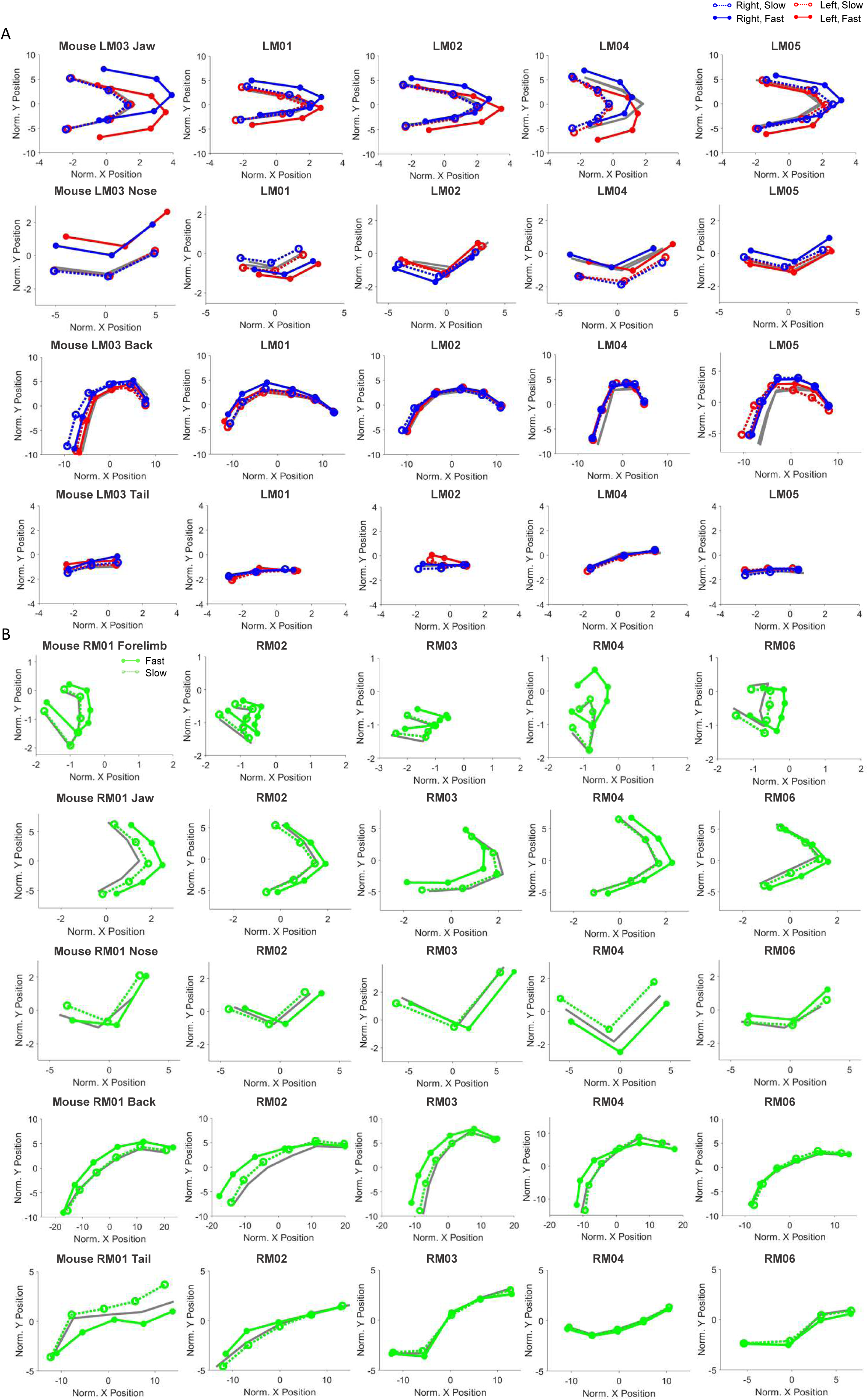
Motor readiness is selectively reflected in task-relevant body features. (A) Body feature readouts in the licking task for all mice. First row: jaw position; second row: nose position; third row: back position; fourth row: tail position. Average body feature position across sessions is calculated at the end of the delay epoch. Blue solid lines denote fast RT lick right trials and red solid lines denote fast RT lick left trials. Blue dashed lines denote slow RT lick right trials and red dashed lines denote slow RT lick left trials. The body feature position at the start of the delay is plotted in gray (baseline). Only correct trials are plotted. Only jaw and nose positions reliably differentiate fast and slow RT trials. (B) Body feature readouts in the reaching task for all mice. First row: forelimb position; second row: jaw position; third row: nose position; fourth row: back position; fifth row: tail position. Green solid lines denote fast RT trials and dashed green lines denote slow RT trials. The body feature position at the start of the delay is plotted in gray. Only forelimb position reliably differentiates fast and slow RT trials.

**Supplemental Figure 4:**
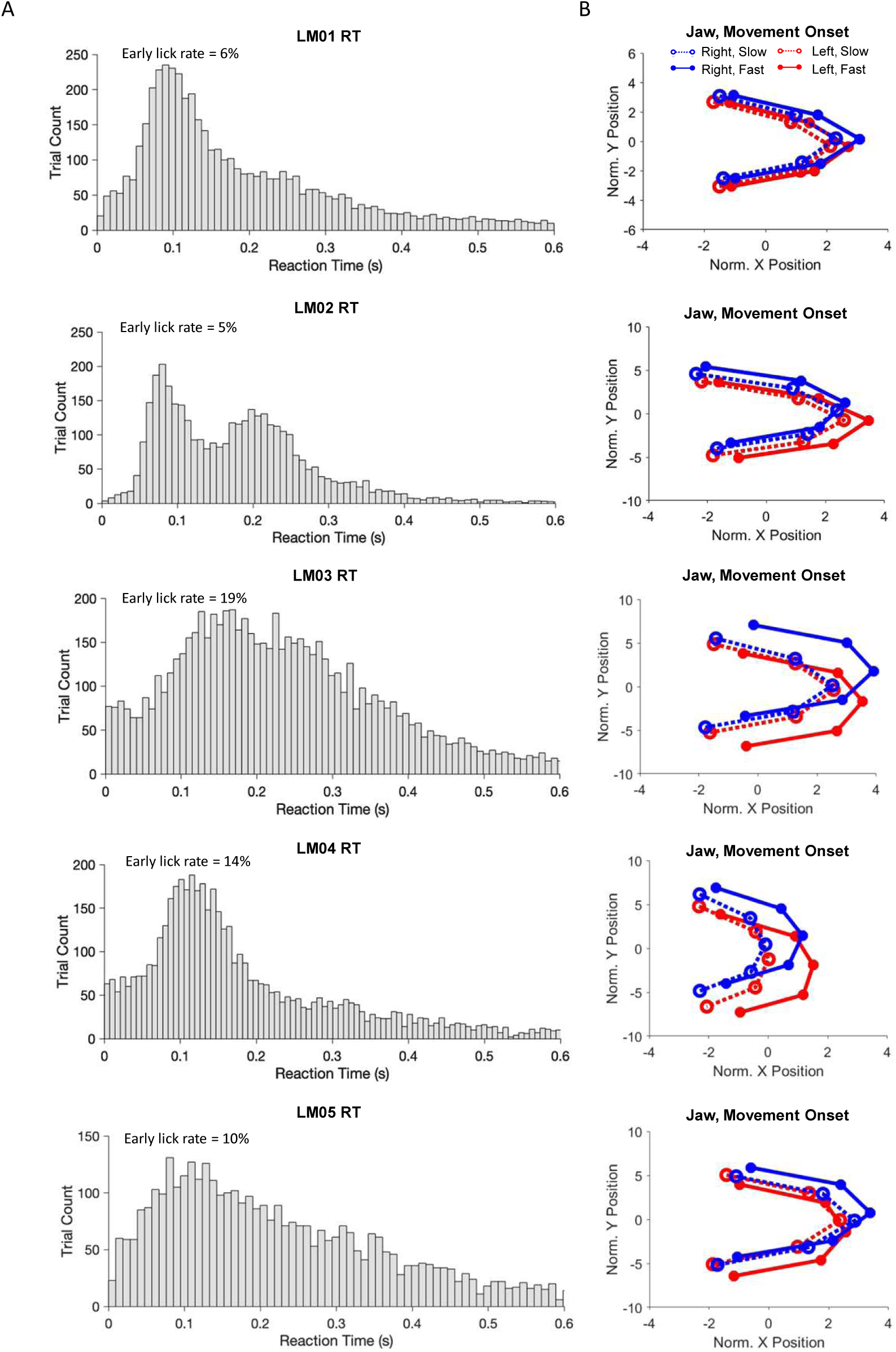
Mice do not show body feature of motor readiness on slow reaction time trials even at movement initiation. (A) Reaction time (RT) distribution for mice in the licking task. RTs are defined as the first video frame in which the tongue is detected from the Go cue onset. On some trials, the tongue is already present upon Go cue arrival. The incident rate is similar to the fraction of trials in which mice licked before the Go cue (“early lick rate”). Early lick rates for each mouse are shown on top. (B) Average jaw position at the moment of movement initiation on slow and fast RT trials. Blue solid lines denote fast RT lick right trials and red solid lines denote fast RT lick left trials. Blue dashed lines denote slow RT lick right trials and red dashed lines denote slow RT lick left trials. On slow RT trials, the jaw position does not achieve the level of displacement seen on fast RT trials, suggesting that body feature displacement reflects motor planning rather than movement initiation.

**Supplemental Figure 5:**
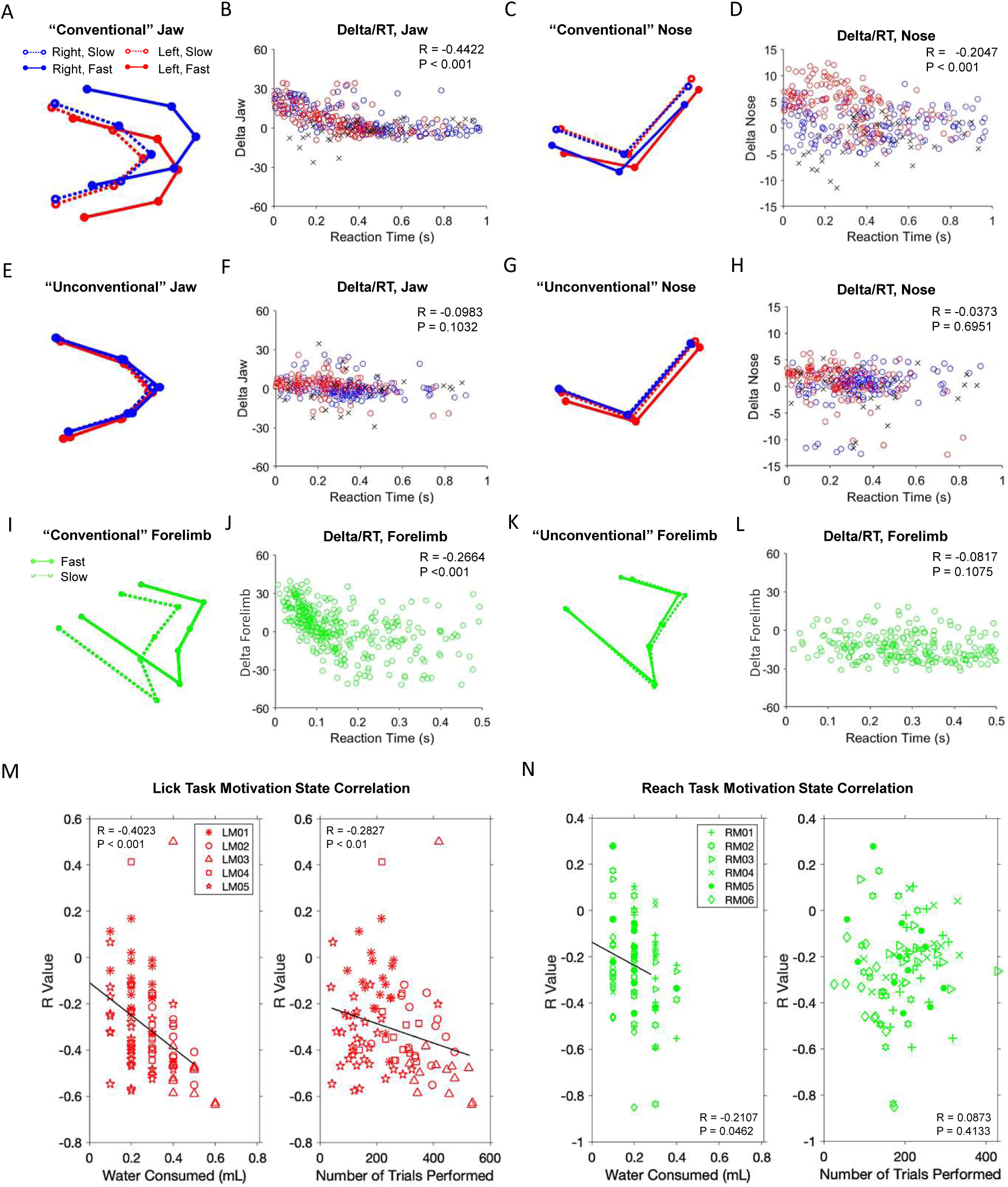
Motor planning is correlated with motivation state. (A) Average jaw positions at the end of the delay epoch in an example “conventional” session. Data from licking task. Blue solid lines denote fast RT lick right trials and red solid lines denote fast RT lick left trials. Blue dashed lines denote slow RT lick right trials and red dashed lines denote slow RT lick left trials. Only correct trials are plotted. Jaw orientation on fast RT trials is much greater than on slow RT trials in conventional sessions. (B) Single trial delta-jaw versus reaction time for the example session shown in (A). Open red circles denote correct lick left trials and open blue circles denote correct lick right trials. Crosses denote incorrect trials. (C) Average nose positions at the end of the delay epoch for the same session shown in (A). (D) Single trial delta-nose versus reaction time for the same session in (A). Delta-nose values are calculated in the same way as delta-jaw values, but in the X dimension. (E) Average jaw positions at the end of the delay epoch in an example “unconventional” session. The amount of jaw orientation on fast RT trials and slow RT trials appears similar in rare unconventional sessions. (F) Single trial delta-jaw versus reaction time for the example session shown in (E). (G)Average nose positions at the end of the delay epoch for the same session shown in (E). (H) Single trial delta-nose versus reaction time for the same session shown in (E). (I) Average forelimb positions at the end of the delay epoch in an example “conventional” session. Data from reaching task. Forelimb displacement is seen on fast RT trials but not on slow RT trials. (J) Single trial delta-forelimb versus reaction time for the example session shown in (I). (K) Average forelimb positions at the end of the delay epoch in an example “unconventional” session. In rare unconventional sessions, there was no difference in forelimb displacement whether RT was fast or slow. (L) Single trial delta-forelimb versus RT for the example session shown in (K). (M)Motivation state correlates with motor readiness in the licking task. Pearson’s correlation (R value) between delta-jaw values and RT is computed across individual trials within each session, providing a behavioral signature of motor planning. Here, session-wise correlation coefficients are compared to two indicators of motivation state: total water consumed and number of trials performed within a session. Mice consumed significantly more water and performed more trials when exhibiting stronger behavioral signatures of motor planning. (N) Motivation state correlates with motor readiness in the reaching task. Same as (M).

**Supplemental Figure 6:**
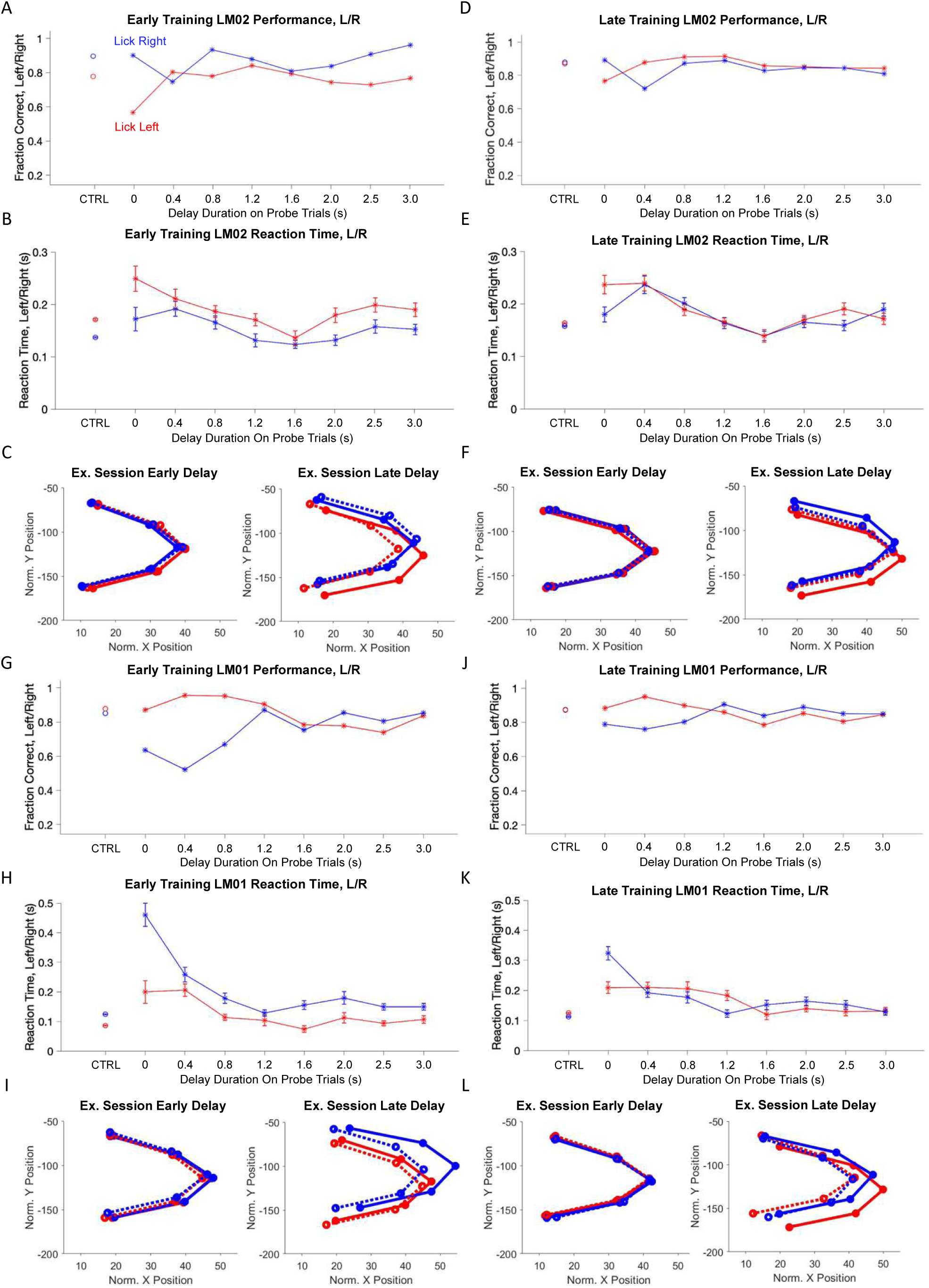
Mice have individual directional biases which can be read out from body features. (A) Task performance (fraction correct) for each trial type for an example mouse that exhibits lick right bias. Data from early training session. Performance is higher on lick right trials (blue) than on lick left trials (red) for control trials and for most probe trial conditions. The performance is especially low for lick left probe trials when the Go cue arrives early (e.g. delay duration, 0 s), indicating that the mouse tends to lick right by default in absence of motor planning. (B) Reaction time (RT) by trial type for the same mouse in (A). RT is consistently faster on lick right trials (blue) compared to lick left trials (red). Error bars indicate the SEM across trials. (C) Jaw position readout of motor readiness for the same mouse in (A). Data from an example early training session. Blue solid lines denote fast RT lick right trials and red solid lines denote fast RT lick left trials. Blue dashed lines denote slow RT lick right trials and red dashed lines denote slow RT lick left trials. The jaw begins in a baseline position that does not differentiate lick direction nor slow vs. fast reaction time trials (left). At the end of the delay epoch, the jaw shows orienting to the right direction selectively on lick right trials, likely reflecting active motor planning (right). In contrast, the jaw remains at the baseline position on lick left trials and does not differentiate slow and fast reaction time trials. This pattern of jaw position is consistent with a default bias to lick left, which is countered by active motor planning to lick right on lick right trials. (D-F) Data from a late training session for the same mouse in (A)-(C). After several training sessions, the lick right bias was reduced. (G-L) Same as (A)-(F) for a mouse that exhibits lick left bias.

**Supplemental Figure 7:**
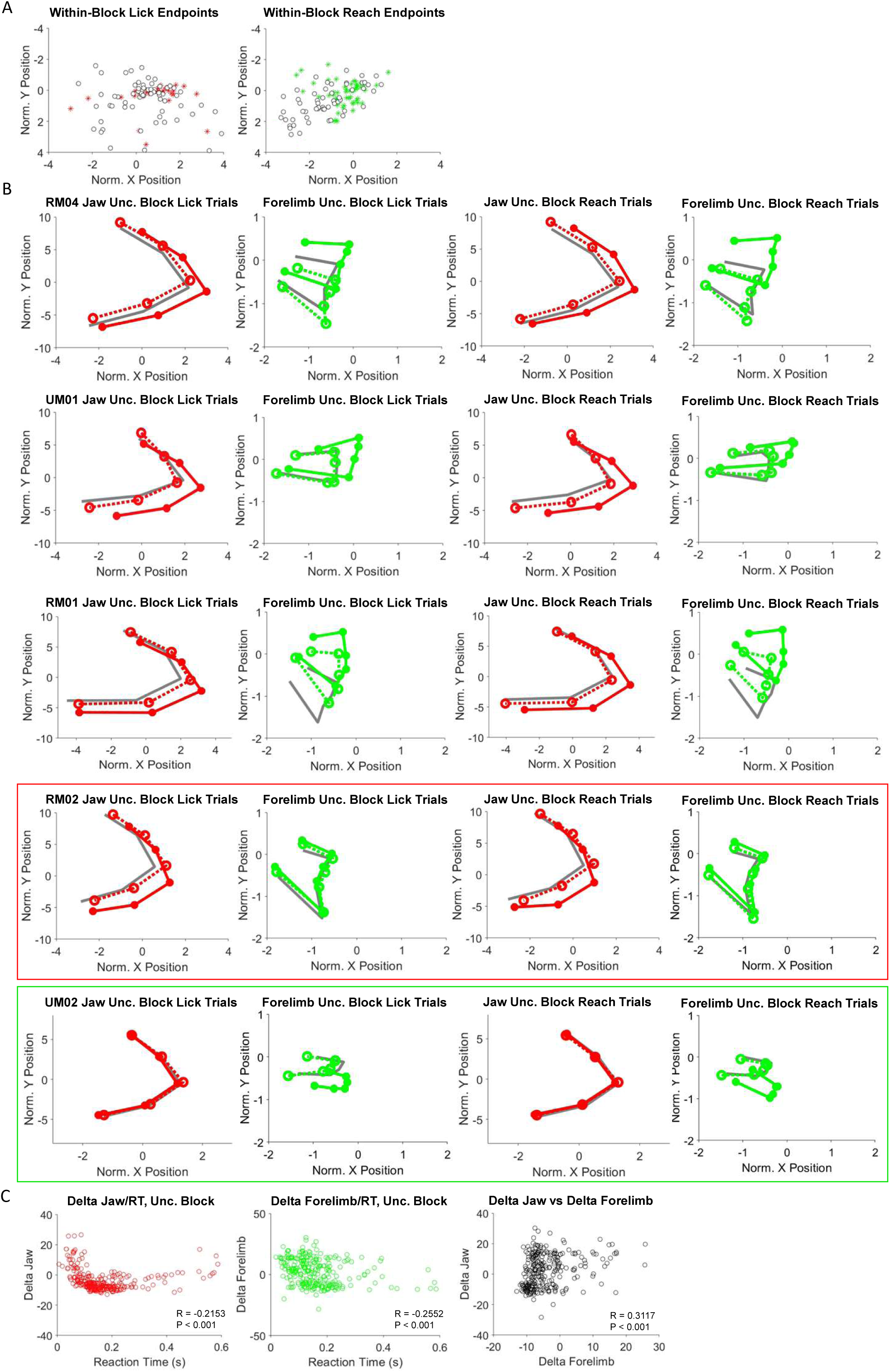
Analysis of movement accuracy and body feature of motor readiness in the uncertain block. (A) *Left*, lick endpoints for the licking trials within the lick block. *Right*, reach endpoints for the reaching trials within the reach block. Data from an example session. Asterisks, fast RT trials; open circles, slow RT trials. On fast RT trials, the movements are more accurate. (B) Average jaw and forelimb positions at the end of the delay epoch in the uncertain block for all mice. Each row shows one mouse. *Left*, trials in which the spout is in position for licking (“lick trials”). *Right*, trials in which the spout is in position for reaching (“reach trials”). A majority of mice show body feature of motor readiness for both licking and reaching in the uncertain block. Red and green boxes, two mice did not demonstrate this simultaneous motor planning. (C) *Left*, single trial delta-jaw versus reaction time for the licking task in the uncertain block. *Middle*, single trial delta-forelimb versus reaction time for the reaching task in the uncertain block. *Right*, single trial delta-jaw versus delta-forelimb in the uncertain block. Both body features were simultaneously displaced on single trials.

